# Decoding Mutually Induced Conformational Changes in Non-Canonical Recognition of U1 SL4 snRNA by ULD of SF3A1 during Early Spliceosome Assembly

**DOI:** 10.64898/2026.03.30.715355

**Authors:** Shri Kant, Atanu Maity, Mashu Bhagat, Savan Maspeddi, Ranjit Prasad Bahadur

**Affiliations:** Computational Structural Biology Laboratory Department of Bioscience and Biotechnology, Indian Institute of Technology Kharagpur, Kharagpur-721302, India; Bioinformatics Center, Department of Bioscience and Biotechnology, Indian Institute of Technology Kharagpur, Kharagpur-721302, India

**Keywords:** Key word: Spliceosome assembly, SF3A1, SL4 of U1 snRNA, ubiquitin-like domain, non-canonical recognition, side chain torsion

## Abstract

A crucial step in early spliceosome assembly is the interaction between splicing factor 3A1 (SF3A1) of U2 snRNP and stem-loop 4 (SL4) of U1 snRNA. This interaction facilitates the spatial alignment of the 5′ and the 3′ splice sites, leading to the formation of the pre-spliceosomal A complex. In this study, we investigate the structural and dynamic basis of non-canonical recognition between ubiquitin-like domain (ULD) of SF3A1 and SL4 of U1 snRNA. Extensive all-atom molecular dynamics simulations reveal a dual recognition mechanism involving sequence-specific interactions mediated by the C-terminal RGGR motif and structural recognition governed by the UUCG tetraloop of SL4 snRNA. The RGGR motif primarily engages the duplex region of the snRNA, whereas the stem-loop nucleotides interact with the globular region of the ULD. Mutations of key residues R788 and R791 result in a significant loss of protein-RNA interactions, as reflected in the reduced binding affinities and altered conformational stability. Nucleotides C6 to C9 in the duplex region, stabilized by strong base-pairing and backbone-mediated interactions with SF3A1, exhibit constrained torsional distributions and minimal sensitivity to mutation. In contrast, nucleotides G10 to C15 in the tetraloop exhibit broader torsional distributions with moderate occupancy, consistent with weaker base-pairing but stronger protein-RNA interactions. Mutations significantly alter torsional distribution, enabling the nucleotides to adopt alternate conformations that preserve interactions with the globular domain of SF3A1. These findings provide a mechanistic insight into non-canonical RNA recognition and highlight the role of coupled sequence and structural determinants in stabilizing early spliceosomal assembly.

## Introduction

Spliceosome is a dynamic large ribonucleoprotein (RNP) machinery responsible for pre-mRNA splicing in eukaryotic cells.^1–3^ Assembly and function of spliceosome is orchestrated by specific protein-RNA interactions. Spliceosome excises non-coding introns and ligates coding exons with single-nucleotide resolution, producing mature messenger RNAs.^4,5^ This intricate machinery assembles during each splicing cycle by undergoing repeated rounds of conformational and compositional remodelling to process a vast array of RNA substrates.^4,6^ A crucial step in early assembly of the spliceosome is the interaction between the splicing factor 3A1 (SF3A1) subunit of the U2 small nuclear ribonucleoprotein (snRNP) and stem-loop 4 (SL4) of the U1 snRNA (Figure 1A).^7,8^ This interaction helps to spatially align the 5′ and 3′ splice sites, leading to the formation of the pre-spliceosomal complex A.^9,10^ Complex A subsequently recruits U4 followed by U6 and U5 tri-snRNP to generate the pre-B complex, which initiates subsequent steps of catalytic activation.^11^ The significance of this early RNA-protein interaction is highlighted by its role in bridging the U1 and U2 snRNPs, thereby establishing the structural foundation of the pre-spliceosome.^1,2,6,7,9^

**Figure 1.**
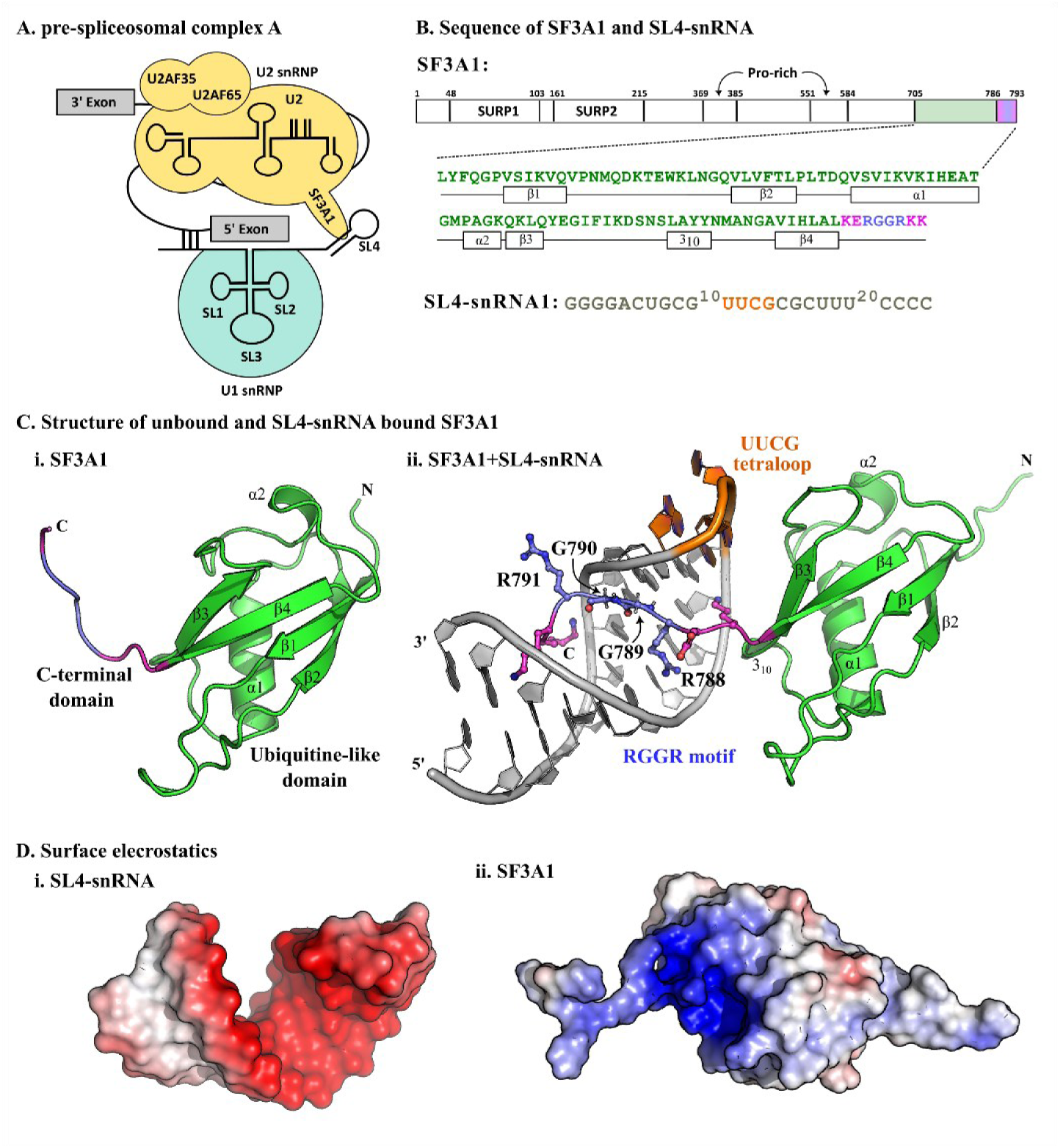
(A) Schematic representation of pre-spliceosomal complex A. (B) Amino acid sequence of the ULD of SF3A1 and nucleotide sequence of SL4-snRNA, highlighting key residues and motifs involved in RNA recognition. (C) Structures of SF3A1 in (i) unbound ULD of SF3A1 (PDB id: 7P08) and in (ii) complex with SL4-snRNA (PDB id: 8ID2), illustrating the interface including the RGGR motif and the UUCG tetraloop. (D) Electrostatic surface potential of (i) SL4-snRNA and (ii) the ULD of SF3A1. Surface potentials are coloured according to the electrostatic scale, where red represents negatively charged regions, and blue represents positively charged regions, indicating complementary charge distributions that facilitate protein-RNA interactions.

SF3A1 function as a part of the SF3A heterotrimer in complex with SF3A2 and SF3A3, and is essential for 17S U2 snRNP maturation and efficient pre-mRNA splicing.^2,10^ SF3A1, a 120 kDa U2 snRNP-specific protein also known as SAP114 or SF3a120, contains two tandem suppressor-of-white-apricot (SURP)^10,12^ domains and a C-terminal ubiquitin-like domain (ULD) with a beta-grasp fold (Figure 1B).^12,13^ The ULD, which lacks classical RNA-binding signatures, mediates direct and specific interaction with SL4 of U1 snRNA.^13,14^ This non-canonical RNA-binding event plays structural and functional roles by tethering U1 and U2 snRNPs, enabling proper recognition and positioning of the splice sites for accurate splicing.^8,15,16^ Although considerable progress has been made in understanding the function of late-stage spliceosomal complex, including U2/U6/U5-mediated catalytic complexes, yet the molecular mechanism underlying early SF3A1-U1 recognition remains elusive.^17,18^ Recent cryo-EM studies in yeast and human reveal interactions between U1 and U2 snRNPs in complex A.^6^ However, the dynamics of SF3A1-SL4 of U1 snRNA interaction including its transition and dissociation are yet to be unravelled.

Emerging evidence suggests that alterations in SF3A1, particularly within the RGG motif of the ULD, lead to spliceosome dysfunction associated with various human diseases. For example, increased SF3A1 due to mis-spliced pre-mRNAs is related to breast cancer.^19^ Rare mutations in SF3A1 expression due to splicing errors affect hematopoietic function in myelodysplastic syndromes.^20^ While SF3A1 has not yet been confirmed as a primary causal gene in ALS, its role in spliceosome integrity and mRNA processing regulation are functionally linked to neurodegeneration.^21^ Affinity and specificity of the interactions between ULD and U1 snRNA is significantly reduced or lost when mutations occur either at the ULD or at the SL4 region.^9,22^ Co-crystal structure of the C-terminal ULD of SF3A1 with the SL4 region of U1 snRNP^9^ reveals that the RGGR motif along with its globular ubiquitin fold interacts with the UUCG tetraloop of snRNA. Another co-crystal structure shows that the stem-loop of snRNA is critical for binding of the single-stranded RNA at the splice sites.^22^ The UUCG tetraloop, a well-known thermodynamically stable motif, caps the SL4 region in U1 snRNA.^23^

Here, we have employed all-atom molecular dynamics (MD) simulations to explore the interface binding dynamics of the ULD of SF3A1in apo and SL4 bound forms. We also study the effect of three critical mutations in ULD affecting the recognition of SL4 of snRNA. Comparative analyses reveal that the mutations induce significant perturbations in conformational stability and interface interactions. The simulations further uncover a dual recognition mechanism: sequence-specific interactions through the RGGR motif and structural recognition mediated by the UUCG tetraloop. Interface Arg residues show higher variation in side chain conformation upon mutation, whereas the neighbouring Lys residues are largely unaffected. The nucleotides at the tetraloop show higher conformational variability upon mutation due to its interaction with the globular ULD domain, whereas the nucleotides at the stem region accommodating C-terminal remains unaffected. These results elucidate the molecular basis of SF3A1-U1snRNA recognition and its implications in spliceosomal assembly.

## Materials and Methods

### Preparation of unbound and bound structures

Structures of ULD domain of SF3A1 in apo form (PDB id: 7P08)^9^ and in complex with U1 SL4 snRNA (PDB id: 8ID2)^22^ were obtained from the Protein Data Bank (PDB).^24^ The asymmetric unit contains two copies of the complex, one of which has three missing residues at the C-terminal of SF3A1. We selected the other complex in the asymmetric unit (protein chain A and RNA chain C) without any missing residues. Additionally, four mutant complexes R788A, R791A, E787A and a dual mutant R788A_R791A were designed using UCSF Chimera.^25^

### Preparation for MD simulation and production run

Each system was solvated in a cubic water box using explicit water molecules (TIP3P)^26^, ensuring a buffer distance of 1 nm between the edges of the box and the system. Sufficient number of counter ions (Na^+^ or Cl^-^) were added to each system to achieve overall charge neutrality. We have used AMBER99SB-ILDN force field^27^ for protein and RNA. All the systems were simulated using GROMACS 2023.^28^ Energy minimization was performed for 50000 steps with a steepest descent (SD) algorithm to remove unfavourable contacts and steric overlaps. Each system was subjected to equilibration run in two phases: NVT equilibration for 500 ps followed by NPT equilibration for 1 ns. Initial velocities were assigned according to the Maxwell distribution^29^ before NVT equilibration to attain 300K temperature. V-rescale thermostat was used for temperature coupling^29^ in NVT and NPT equilibration steps. Parrinello-Rahman barostat^30^ was used for pressure coupling to maintain a pressure of 1 atmosphere in NPT equilibration. Production dynamics of 500 ns was carried out for each system after the equilibration. Timestep of 1 fs was used for NVT equilibration stage. The LINCS algorithm^31^ was used to constrain all bonds, including hydrogen bonds (H-bond) during NPT equilibration and production dynamics to allow 2 fs timestep. A cut-off distance of 1 nm was used for the calculation of short-range electrostatics and van der Waals interactions. Long-range electrostatic interactions were computed using the Particle-Mesh Ewald (PME)^32^ summation method with a periodic boundary condition. Coordinates and velocities were saved at an interval of 10 fs during the production dynamics. All simulations were duplicated by reassigning the initial velocities before NVT equilibration.

### Simulation convergence and trajectory analysis

Trajectories obtained from the simulations were analysed to extract several structural parameters and thermodynamic quantities. Root mean square deviation (RMSD), radius of gyration (Rg) and root mean squared fluctuation (RMSF) were calculated using different analysis modules of GROMACS to evaluate the conformational changes. Clustering was performed using *gmx_cluster* to derive most populated clusters and their representative structures for each system. Intermolecular H-bonds were computed using cpptraj^33^, while the occupancy of H-bonds was determined through an in-house Python script. Additional intermolecular interactions and structural parameters were computed using in-house Python scripts.

### Free energy of binding

The binding affinities of both wild-type and mutant complexes were calculated by Molecular Mechanics Poisson-Boltzmann Surface Area (MM-PBSA) method using gmx MMPBSA.^34^ The free energy of binding (ΔG_binding_) was calculated using Equation 1, where TΔS represents the entropic component. The molecular mechanics potential energy (ΔE_MM_) was calculated using van der Waals interactions and electrostatic interactions was calculated using Poisson-Boltzmann (PB) equation. Solvation free energy (ΔG_solvation_) was determined from both polar and non-polar parts using solvent-accessible surface area (SASA). The default probe radius of 1.4 Å was used for SASA calculation. The dielectric constants for the solute and the solvent were assigned to four and 80, respectively. Last 100 ns of the simulation trajectories were used to compute all relevant energy components.

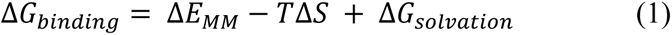

### Calculation of protein side chain rotamers

Protein backbone and side chain conformations adopt distinct rotameric states that are critical for molecular interaction and recognition. To investigate side chain rotamer preferences, we analysed protein side chain conformations across six systems: WT_unbound, WT, R788A, R791A, R788A_R791A and E787A. side chain torsion angles (χ) were computed for 18 selected amino acids for each system on a frame-by-frame basis using the mdtraj.^35^ The analysis was performed over the last 400 ns of each trajectory, excluding the initial 100 ns equilibration phase. The χ torsion angles were defined according to standard geometric conventions. Specifically, χ_1_ represents rotation around the C_α_-C_β_ bond and is defined as the dihedral angle formed by the N-C_α_-C_β_-X_γ_ atoms. χ_2_ corresponds to rotation around the C_β_-X_γ_ bond (C_α_-C_β_-X_γ_-X_δ_), χ_3_ around the X_γ_-X_δ_ bond (C_β_-X_γ_-X_δ_-X_ε_), χ_4_ aound the X_δ_-X_ε_ bond (X_γ_-X_δ_-X_ε_-X_ζ_), and χ_5_ around the X_ε_-X_ζ_ bond (X_δ_-X_ε_-X_ζ_-X_nh1_), where X denotes the appropriate side chain atom at each position. The χ-angle distributions were categorised into three canonical rotameric states including gauche⁺ (*p*; 30° < χ < 90°), trans (*t*; 150° < χ < 150°) and gauche⁻ (*m*; −90° < χ < −30°); and three non-canonical off-rotameric states including (*T*; −30° < χ < 30°), (*P*; 90° < χ < 150°) and (*M*; −150° < χ < −90°). For comparison across all systems, polar histograms were generated to visualise rotamer populations and assess differences in side chain conformational preferences among the six trajectories.

### Calculation of RNA conformation

Structural parameters of RNA were extracted from the MD trajectories using Barnaba^36^ package. The trajectories were analysed to characterise changes in the RNA backbone conformations and ribose sugar pucker by computing the six canonical backbone torsion angles (α, β, γ, δ, ε, ζ) along with sugar torsion angles (V0 to V4), and nucleobase glycosidic torsion angles (χ). The backbone dihedral angles were defined using standard atom selections. Generally, α is defined by O3′(i-1)- P(i)-O5′(i)-C5′(i), β by P(i)-O5′(i)-C5′(i)-C4′(i), γ by O5′(i)-C5′(i)-C4′(i)-C3′(i), δ by C5′(i)-C4′(i)-C3′(i)-O3′(i), ε by C4′(i)-C3′(i)-O3′(i)-P(i+1) and ζ by C3′(i)-O3′(i)-P(i+1)-O5′(i+1). The two pseudo-torsion angles were calculated as η (defined by C4′(i−1)-P(i)-C4′(i)-P(i+1)) and θ (defined by P(i)-C4′(i)-P(i+1)-C4′(i+1)), providing a reduced representation of the backbone geometry of RNA. The glycosidic torsion angle χ was computed to describe base orientation relative to the sugar moiety by using the atom sequence O4′-C1′-N9-C4 for purines and O4′-C1′-N1-C2 for pyrimidines. Together, these parameters provide a comprehensive description of RNA backbone flexibility, sugar pucker dynamics and base orientation changes across all simulated systems.

## Results

### Structures of unbound SF3A1 and its complex with SL4 RNA

Three-dimensional structures of free SF3A1 and its bound form with SL4-snRNA are shown in Figure 1C. The ULD of SF3A1 features a two-layer α-β architecture with a β1-β2-α1-β3-α2-β4 topology (Figure 1C(i)). An additional 3_10_ helix (residues 752 to 754) is present in the complex structure, which joins α1 and β3 (Figure 1C(ii)). The UUCG tetraloop interacting with SF3A1 is highlighted in orange. Interface residues and nucleotides identified in the native complex (PDB id: 8ID2) are provided in Table 1.

**Table 1.** List of interface residues and nucleotides identified from the crystal structure of bound SF3A1 with SL4-snRNA.

| System | Method | Interface residues |  |  |  |
| --- | --- | --- | --- | --- | --- |
|  |  | SF3A1 protein |  | SL4-snRNA |  |
|  |  | Count | Residue's list | Count | Nucleotide list |
| 8ID2 | SASA-based<sup>a</sup><br>(NACCESS) | 14 | Gly753, Lys754, Lys756,<br>**Phe763**, Lys765, Asp766,<br>Lys786, Glu787, **Arg788**,<br>**Gly789**, **Gly790**, Arg791,<br>Lys792, Lys793 | 21 | G1, G2, G3, G4, C6, U7, G8,<br>C9, G10, ***U11***, ***U12***, ***C13***,<br>C15, G16, C17, U18, U19,<br>U20, C21, C22, C23 |
| 8ID2 | Distance-based<sup>b</sup><br>(cut-off 3.5 Å) | 12 | Lys754, Lys756, **Phe763**,<br>Lys765, Lys786, Glu787,<br>**Arg788**, **Gly789**, **Gly790**,<br>Arg791, Lys792, Lys793 | 12 | G2, G3, U7, G8, C9, G10,<br>***U11***, C13, C15, G16, C17,<br>U18 |

Interface residues reported as critical binding residues are marked in bold (N. Nameki et al. (2023)).
Interface nucleotides from tetraloops (UUCG) are marked in bold italics (N. Nameki et al. (2023)).
<sup>a</sup>NACCESS is used to calculate SASA using default parameters. Residues whose atoms lost SASA upon complexation are identified as interface residues.
<sup>b</sup>Euclidean distance between each pair of protein and RNA atoms is calculated from their coordinates in PDB. Any atom pairs within a cut-off distance of 3.5 Å are considered as interacting.

Eight-residues long C-terminal tail of SF3A1 (KERGGRKK) is disordered, which extensively interact with the major groove of the RNA (Figure 1C(ii)). Electrostatic potential surfaces illustrate continuous patch of similar electrostatic potential, where negative potential (red surface in Figure 1D(i)) arises due to an abundance of negatively charged phosphate groups at the RNA backbone. In contrast, region of positive potential (blue surface in Figure 1D(ii)) corresponds to positively charged amino acid residues of SF3A1.

### Structural flexibility of the C-terminal domain of SF3A1 in RNA recognition

Convergence of the 500 ns long trajectories is assessed by calculating root mean square average correlation (RAC), which is computed using CPPTRAJ^33^ for all trajectories. RAC is an essential metric for evaluating the convergence of trajectories. Root mean square deviation (RMSD) typically shows how much a structure deviates from a reference and can be subjective in interpreting equilibration. Whereas, RAC offers a more robust approach and quantifies how closely individual frame in a trajectory aligns with the average structure of all the frames. To calculate RAC, each simulation frame is first aligned (RMS-fit) to the average structure derived from all frames in the trajectory. The average RMSD is computed and monitored as a function of time throughout the trajectory. As simulation converges, RAC values should decrease and eventually level off, indicating that further sampling does not significantly alter the ensemble structure. The RAC profiles display steady decreasing slopes beyond 150 ns, approaching zero near 500 ns for all the systems and their replicates (Figure S1). Notably, RAC rapidly decline after 200 ns with an average all-atom RMSD below 0.5 nm. This consistent pattern indicates that each simulation has reached equilibrium with internal structural fluctuations stabilizing over time.

Average backbone RMSD of WT_unbound is significantly higher than that of WT (Figure 2A). The four mutant complexes also exhibit lower RMSD (average is less than 0.3 nm) than WT_unbound. This indicates that higher fluctuations in unbound SF3A1 is reduced upon binding U1 snRNA. WT_unbound shows the minimum Rg with an average of 1.35 nm. In contrast, WT exhibits consistently higher Rg throughout the simulation (Figure 2B). Among the mutants, most of them show higher Rg than WT_unbound. Interestingly, R788A_R791A shows Rg in between WT_unbound and WT. The difference in the range and variation of RMSD and Rg profiles is further investigated by looking into the residue level fluctuation of the SF3A1 residues. Simulated trajectories of the replicates show similar trends (Figure S2).

**Figure 2.**
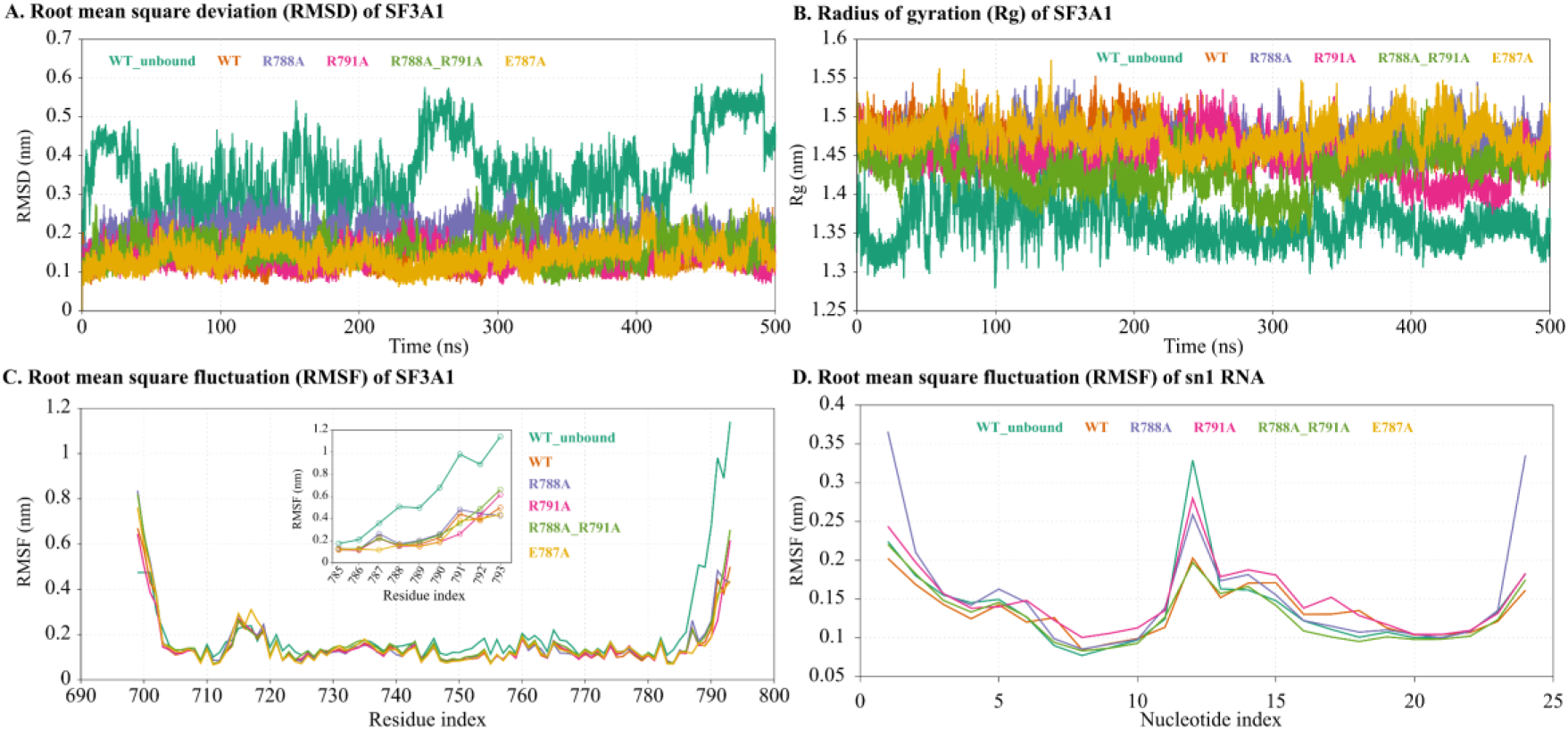
(A) Root mean square deviation of SF3A1, (B) radius of gyration (Rg) of SF3A1. (C) root-mean-square fluctuation (RMSF) of SF3A1, and (D) RMSF of the of SL4-snRNA.

### Conformational flexibility in RNA recognition

Conformational flexibility of SF3A1-SL4RNA recognition is evaluated using RMSF (Figure 2C). WT_unbound shows higher RMSF than WT and other mutants, especially in the C-terminal region of SF3A1. The RMSF of C-terminal residues in WT and mutants are approximately 0.3 nm lower than that in the unbound SF3A1 (Figure 2C inset). This suggests that the C-terminal region of the ULD is structurally more flexible in unbound SF3A1 compared to SL4-snRNA complex. Moreover, the extreme C-termini disordered residues R791, K792 and K793 have varying RMSF in different mutants. In the unbound state, these residues show random motion in extended conformation and often interact with the globular region of SF3A1. This leads to higher RMSD and lower Rg of unbound SF3A1 compared to its bound states.

In RNA, U12 and C13 of the UUCG tetraloop exhibit higher fluctuations than the terminal nucleotides, which show increased variability at both 52 and 32 ends across all the systems except R788A. U12 in WT exhibits higher fluctuations relative to other mutants. Conversely, 52 G1 and 32 C24 exhibit greater fluctuations in R791A relative to WT. Among all the mutants, the nucleotides in R788A exhibit the least fluctuation.

### Analysis of intermolecular interactions

We have analysed H-bonds, stacking interactions, salt bridges and hydrophobic interactions between SF3A1 and SL4-snRNA for the representative complex of the most populated cluster. The amino acids involved in different types of interactions with SL4-snRNA are listed in Table 2.

**Table 2.** List of residues involved in most occupied intermolecular interactions in WT and mutant complexes.

| System | Type of Intermolecular interactions |  |  |  |
| --- | --- | --- | --- | --- |
|  | H-bonds<sup>a</sup> | Stacking<sup>b</sup> | Salt-bridge<sup>c</sup> | Hydrophobic<sup>d</sup> |
| WT | Lys756, Lys786, Glu787, Arg788, Gly789, Lys792 | Arg788, Arg791 | Lys754, Arg791, Lys793 | Phe763 |
| R788A | Lys756, Ile764, Lys786, Gly789, Arg791, Lys792, Lys793 | Arg791 | Lys717, Arg791 | Pro751, Ala752, Phe763, Ile764, Ala788 |
| R791A | Lys756, Glu787, Arg788, Gly789, Lys792 | NA | Lys793 | Phe763, Ala791 |
| R788A_R791A | Lys756, Glu787, Ala791, Lys792 | NA | Lys754, Lys765, Lys793 | Phe763, Ala788, Ala791 |

<sup>a</sup>H-bonds are calculated using HBPLUS with default parameters.
<sup>b</sup>Stacking interactions are defined as p-p and cation-p interactions formed by aromatic and charged residues with nucleotide bases with a cut-off distance of 6.5 Å and dihedral angle less than 30°.
<sup>c</sup>Salt-bridges are identified as interactions within 4.0 Å distance between side chain N-atoms of positively charged residues and negatively charged phosphates of nucleotides.
<sup>d</sup>Hydrophobic interactions are calculated based on pairwise interatomic interactions between atoms of hydrophobic residues within a cut-off distance of 4.5 Å.

In WT, Arg788 and Arg791 maintain native stacking and salt-bridge interactions. Their mutations lead to significant reorganization of intermolecular interactions. In both the single mutants, the loss of key Arg residues disrupts stacking interactions and reduces salt-bridge interactions. This loss is partially compensated by a newly formed intermolecular H-bonds involving Ile764 and Lys793 in R788A. In the double mutant, stacking interactions and salt bridges are completely lost. Instead, hydrophobic interactions increased due to the mutation to Ala.

### Free energy of binding and residue decomposition

Binding free energy (ΔG) reveals a clear trend of reduced RNA binding affinity in all the mutants, except E787A (Table 3). The WT shows the strongest binding affinity with ΔG of −112.29 kcal/mol. Among the mutants, R791A exhibits a substantial loss of binding affinity (ΔG = −56.39 kcal/mol), followed by R788A_R791A (ΔG = −62.26 kcal/mol) and R788A (ΔG = −77.93 kcal/mol). The sum of electrostatic interaction energy and polar solvation energy (ΔE_elec_+ΔE_polar_) is a measure of energetic gain or loss when the solvated protein and RNA are complexed together. These two energy components usually carry opposite signs, which implies that if combining the two produces favourable electrostatic interaction there will be a loss of polar solvation energy. Their sum represents the overall polar interaction including solvation upon binding. In all the mutants, the van der Waals interaction energy is significantly lower than the WT. Sum of ΔE_elec_ and ΔE_polar_ also shows a significant drop in three mutants, even resulting in positive values for R788A and R788A_R791A. This results in a significant loss of binding affinity for these mutants. In contrast, the loss in van der Waals interaction energy has been compensated by an increase of ΔE_elec_+ΔE_polar_ in E787A, resulting in similar binding affinity to that of WT.

**Table 3.** Binding free energy components of WT complex and mutants.

| Systems | Energy components (kcal/mol) |  |  |  |  |  |
| --- | --- | --- | --- | --- | --- | --- |
| | $\Delta E_{\text{vdw}}$ | $\Delta E_{\text{elec}}$ | $\Delta E_{\text{polar}}$ | $\Delta E_{\text{non-polar}}$ | $\Delta E_{\text{elec}} + \Delta E_{\text{polar}}$ | $\Delta G$ |
| WT | -75.75<br>$\pm 12.52$ | -3555.16<br>$\pm 128.20$ | 3528.5<br>$\pm 125.31$ | -9.89<br>$\pm 1.05$ | -26.66 | -112.29<br>$\pm 14.99$ |
| R788A | -69.81<br>$\pm 9.96$ | -2940.05<br>$\pm 115.82$ | 2941.59<br>$\pm 116.07$ | -9.67<br>$\pm 0.88$ | 1.54 | -77.93<br>$\pm 12.83$ |
| R791A | -42<br>$\pm 31.93$ | -1988.93<br>$\pm 117.79$ | 1980.11<br>$\pm 117.51$ | -5.57<br>$\pm 4.18$ | -8.82 | -56.39<br>$\pm 19.60$ |
| R788A_R791A | -57.63<br>$\pm 8.62$ | -2218.17<br>$\pm 110.29$ | 2221.26<br>$\pm 109.34$ | -7.72<br>$\pm 0.81$ | 3.09 | -62.26<br>$\pm 11.31$ |
| E787A | -66.95<br>$\pm 8.69$ | -4047.11<br>$\pm 128.13$ | 4012.15<br>$\pm 124.32$ | -9.22<br>$\pm 0.7$ | -34.96 | -111.13<br>$\pm 13.31$ |

$\Delta E_{\text{vdw}}$ , $\Delta E_{\text{elec}}$ , $\Delta E_{\text{polar}}$ and $\Delta E_{\text{non-polar}}$ are the energy components for van der Waals, electrostatic, polar and nonpolar solvation energies and $\Delta G$ represents the total binding free energy.

Further, residue-wise decomposition of binding free energy shows distinct energetics in WT and other mutants (Figure 3A). The energetic contributions of amino acids and nucleotides in the binding pocket are plotted for all the systems. Residues with contribution less than −2 kcal/mol (favourable) and more than 2 kcal/mol (unfavourable) are shown in figure 3A(i). The contributions of nucleotides higher than 1.5 kcal/mol and lower than −1.5 kcal/mol are shown in figure 3A(ii). Residues R788 and R791 in WT show relatively higher contribution with −12.1 kcal/mol and −15.1 kcal/mol, respectively. In addition, residues K754, K765 and K792 show significant favourable contribution. Interestingly, residues K786 and E787 show strong unfavourable contribution to overall binding energy. Upon mutation, the contribution of R788 decreases below −2 kcal/mol along with a drop in the contribution of R791 (−15.1 kcal/mol to −9.5 kcal/mol). Similarly, mutation of R791 causes a drop in its contribution below −2 kcal/mol accompanied by significant loss in contribution of R788 (−12.1 kcal/mol to −6.8 kcal/mol). These mutations also reduce the energy contribution of two Gly (G789 and G790) present in-between R788 and R791. When the above two arginine’s are simultaneously mutated in R788A_R791A, contributions of both of them drops below −2 kcal/mol along with the reduction of contribution of G789 and G790 as well. In E787A, the contribution of all these residues are similar to the WT system without affecting the overall binding affinity. Among the nucleotides of SL4-snRNA, G10, C13 and C15 show the highest favourable contribution to binding free energy in WT (Figure 3A(ii)). Upon mutation of R788 and R791, contributions of most of the nucleotides decreases. In R788A, U7 and G8 even contribute in unfavourable interactions. Compared to the stem region, the two ends of the RNA are involved in weaker interactions with the C-terminal residues of SF3A1. Mutation of charged amino acids leads to further loss of these weaker interactions, leading to reduced binding affinity. E787A mutation maintains most of the contributions and hence the binding affinity remains almost unaffected.

**Figure 3.**
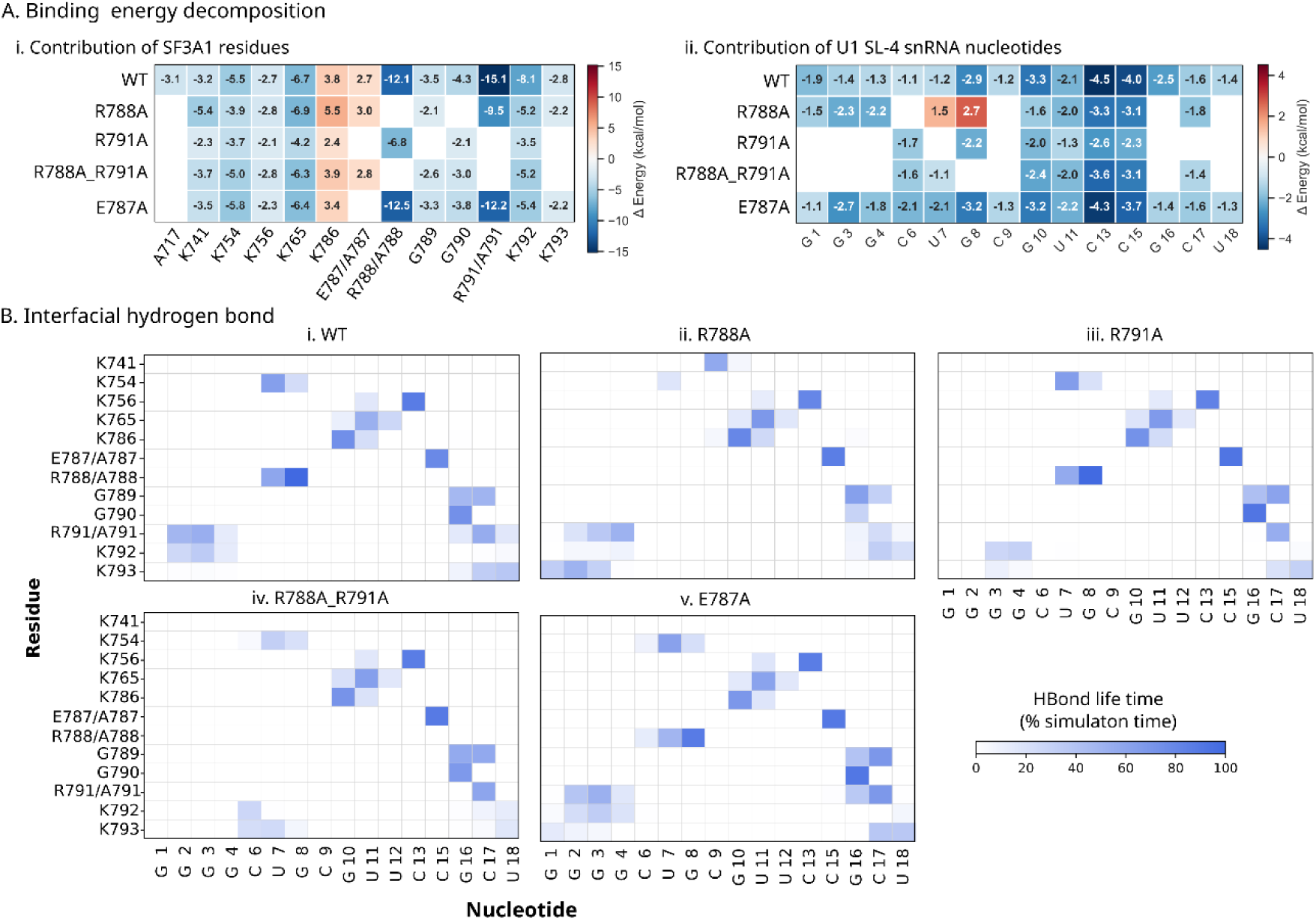
Binding energy decomposition and interface H-bonds of the SF3A1–SL4-snRNA complexes. (A) Per-residue and per-nucleotide decomposition of binding energy. Contribution of individual residues and nucleotides to the total binding free energy of the ULD of SF3A1 (i) and (ii) SL4-snRNA, respectively. Colour scales represent the change in binding free energy (ΔE, kcal/mol), where negative values (blue) indicate favourable contributions and positive values (red) indicate unfavourable contributions. (B) Distribution of interface h-bonds between SF3A1 residues and SL4-snRNA nucleotides. Occupancy of H-bonds over the simulation time are shown for (i) wild type (WT), (ii) R788A, (iii) R791A, (iv) R788A_R791A and (v) E787A. Colour intensity corresponds to hydrogen-bond lifetime (percentage of simulation time), highlighting key interactions that stabilize the protein-RNA complex.

### Dynamics of hydrogen bonds

The heatmap in figure 3B shows the lifetime of H-bonds between the interface amino acid and nucleotides in terms of percent of simulation time. The residue R788 makes H-bond with U7 and G8, whereas R791 makes H-bonds with G2, G3 and G4 in the WT complex (Figure 3B(i)). The mutation of R788 to Ala in R788A and R788A_R791A leads to the loss of these H-bonds with U7 and G8. Due to R791A mutation in R791A and R788A_R791A, the H-bonds involving R791 and G2, G3 and G4 is lost. The H-bond lifetime of residues K741, K754, K756, K765 and K786 with C9-C13 segment remains similar across the systems. Interestingly, the H-bond between E787 and C15 remains unaffected in E787A, highlighting the presence of a backbone mediated H-bond. Residues G789 and G790 forms backbone mediated H-bonds with G16 and C17, which remains intact across different systems with minor variation in their life time.

The long side chain interface Arg and Lys form multiple H-bonds mediated by their backbone and side chain with the backbone and bases of interface nucleotides. The variations in H-bond conformations of the interface amino acids are shown as heatmaps in Figure 4. Figure 4 describes the backbone mediated H-bonds. Residues E787, G789, G790 and R791 has significant lifetime of backbone-mediated H-bonds with RNA. In mutants R788A, R791A and R788A_R7891A, the lifetime of h-bond remains similar. G789 and G790 show high h-bond lifetime along with E787, whereas K792 engaged in some backbone mediated h-bonds in the double mutant. Interestingly, the backbone mediated h-bond is lost in E787A, which is compensated by new h-bonds between backbone of R788 and K792 with the RNA.

**Figure 4.**
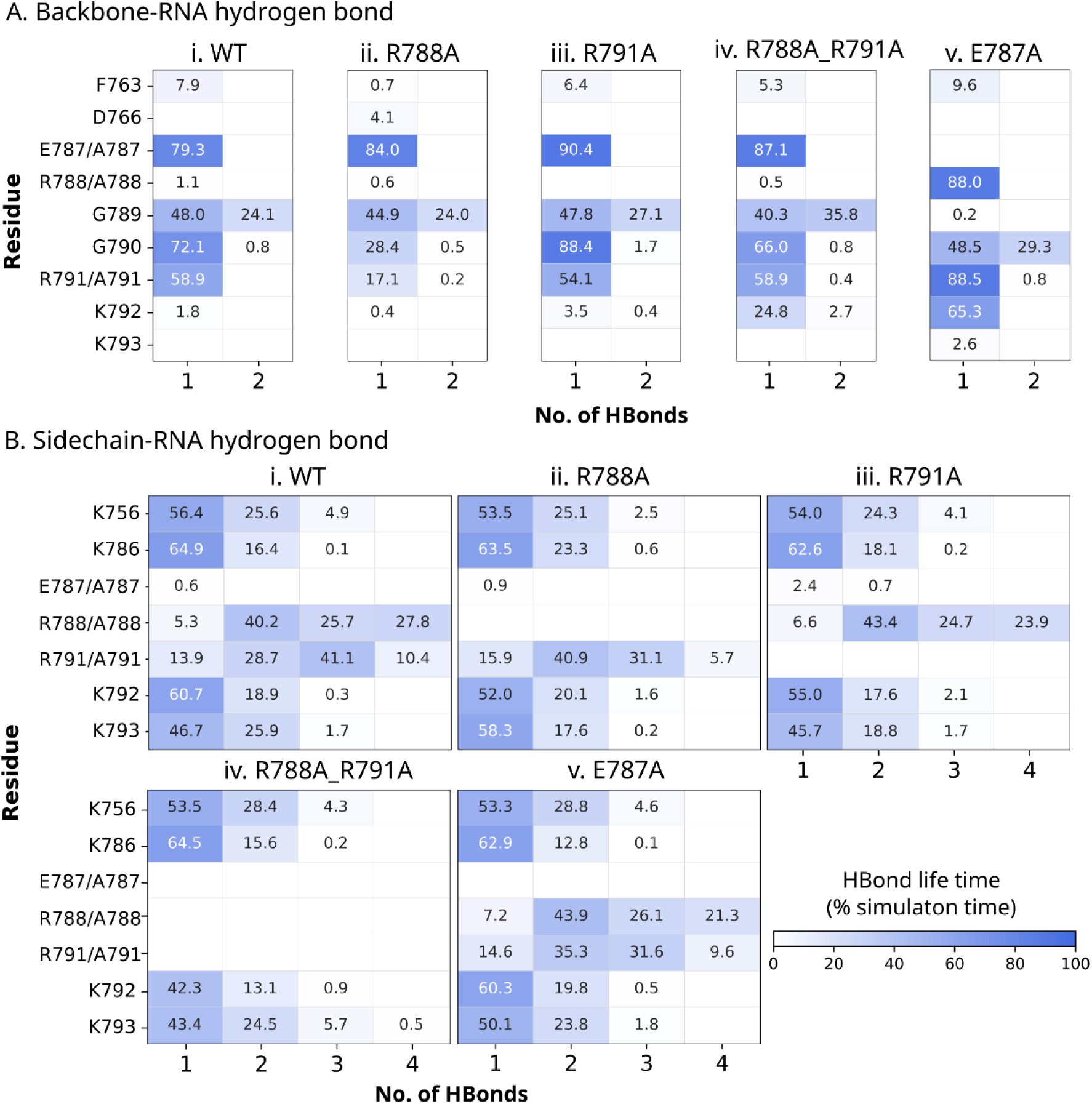
Analysis of backbone and side chain mediated H-bonds in SF3A1–SL4-snRNA complexes. Heatmaps represent the lifetime of H-bonds (percentage of simulation time over 500 ns) between SF3A1 residues and SL4-snRNA. (A) H-bonds mediated by the amino acids backbone of ULD and RNA in (i) wild type (WT), (ii) R788A, (iii) R791A, (iv)R788A_R791A and (v) E787A mutant. (B) H-bonds mediated by side chain of ULD and RNA in (i) WT, (ii) R788A, (iii) R791A, (iv) R788A_R791A, and (v) E787A.Colour intensity reflects hydrogen-bond occupancy, highlighting key residue-nucleotide interactions and their modulation upon mutation.

The variation in side chain-mediated H-bonds is shown in Figure 4B. Despite its anionic side chain, E787 does not form any significant side chain-mediated H-bond. The lysine residues (K756, K786, K792 and K793) form single H-bond with their side chain in 50% of the conformations sampled. In ∼20% of the conformations, they form two side chain-mediated H-bonds. Three side chain-mediated H-bonds are rare due to the presence of only one nitrogen atom (NZ) and three polar hydrogens attached to it. Along with five polar hydrogens, Arg contains three nitrogen atoms (NE, NH1 and NH2), facilitating the formation of multiple H-bonds with different partners simultaneously. Conformations with two H-bonds have the highest lifetime (∼ 40%). The simultaneous presence of three and four H-bonds is also abundant with lifetime of ∼25%-30% of simulation time.

### side chain rotamer dynamics in wild-type and mutant complexes

We investigate the role of side chain conformational dynamics of SF3A1 in recognising SL4 RNA in terms of the side chain torsions of interacting residues. The distributions of side chain torsion of five interface residues are shown in Figures 5-7, and their values are provided in Table S1 and S2. In the complex systems (WT and four mutants), torsion angles are calculated on a set of selected frames in which amino acids are involved in interface H-bonds. We have calculated the possible unique H-bonds (at the level of donor, hydrogen and acceptor atom) involving each of the five amino acids and calculated their lifetime. The H-bond with the longest lifetime for the selected amino acid is considered and the corresponding frames are extracted. All possible side chain torsion angles of the residues are calculated and their distributions are compared with WT_unbound (Figure 5-7). All side chain conformations are considered from the last 400 ns in case of WT_unbound. This ensures comparison of the side chain conformation in unbound state with those involved in a significantly strong and specific H-bond interactions in the complexes.

**Figure 5.**
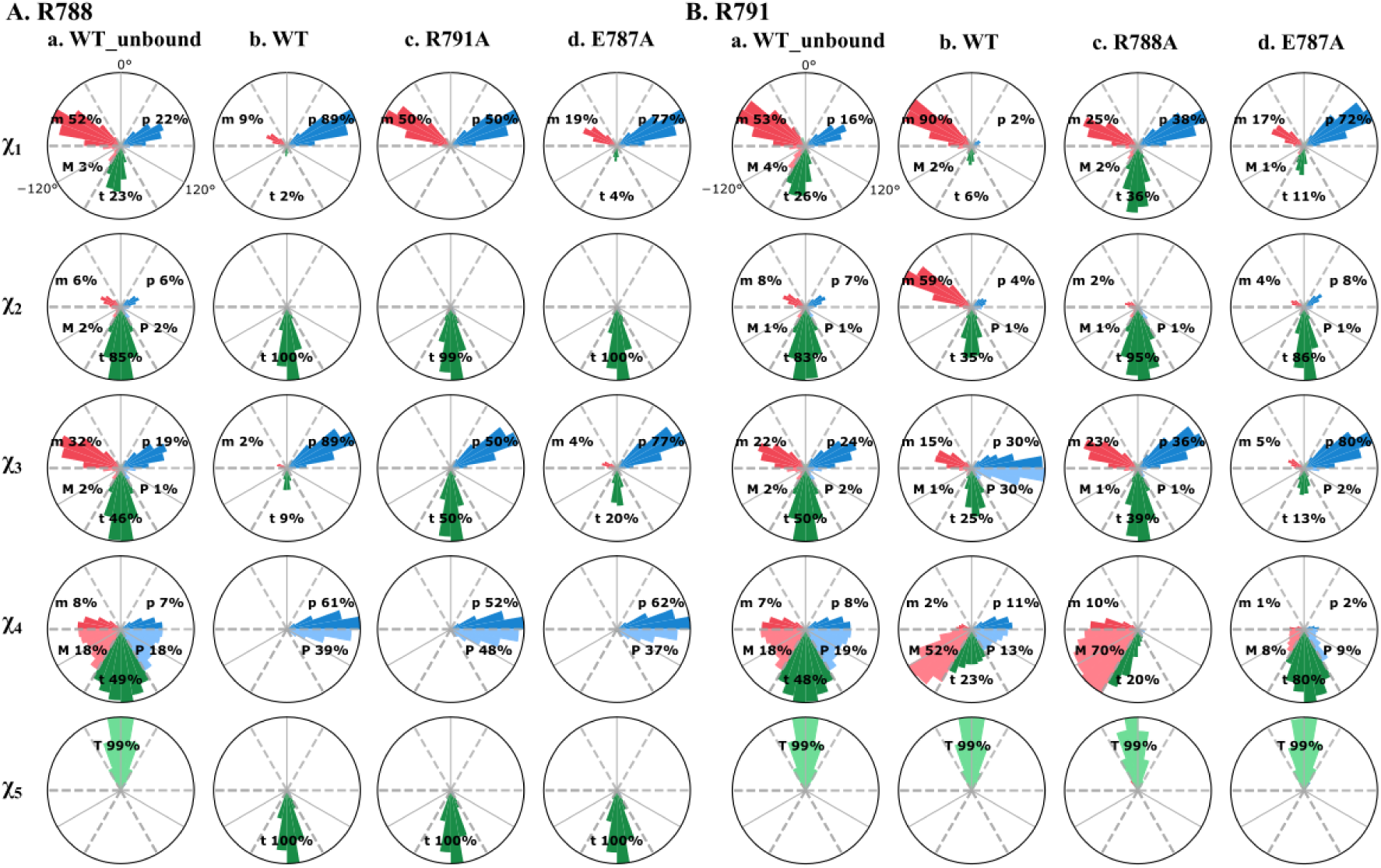
Population distribution of side chain rotamers of Arg788 and Arg791. Polar histogram represents the distributions of side chain dihedral angles (χ₁-χ₅) for (A) Arg788 and (B) Arg791 across different systems: (a) WT_unbound, (b) WT, (c) single mutants (R791A or R788A), and (d) E787A mutant. The distributions are shown as percentage populations of rotameric states, highlighting preferred conformations sampled during the simulations. Comparison between bound and unbound states, as well as across mutants, reveals mutation-induced shifts in rotamer preferences and side chain flexibility.

**Figure 6.**
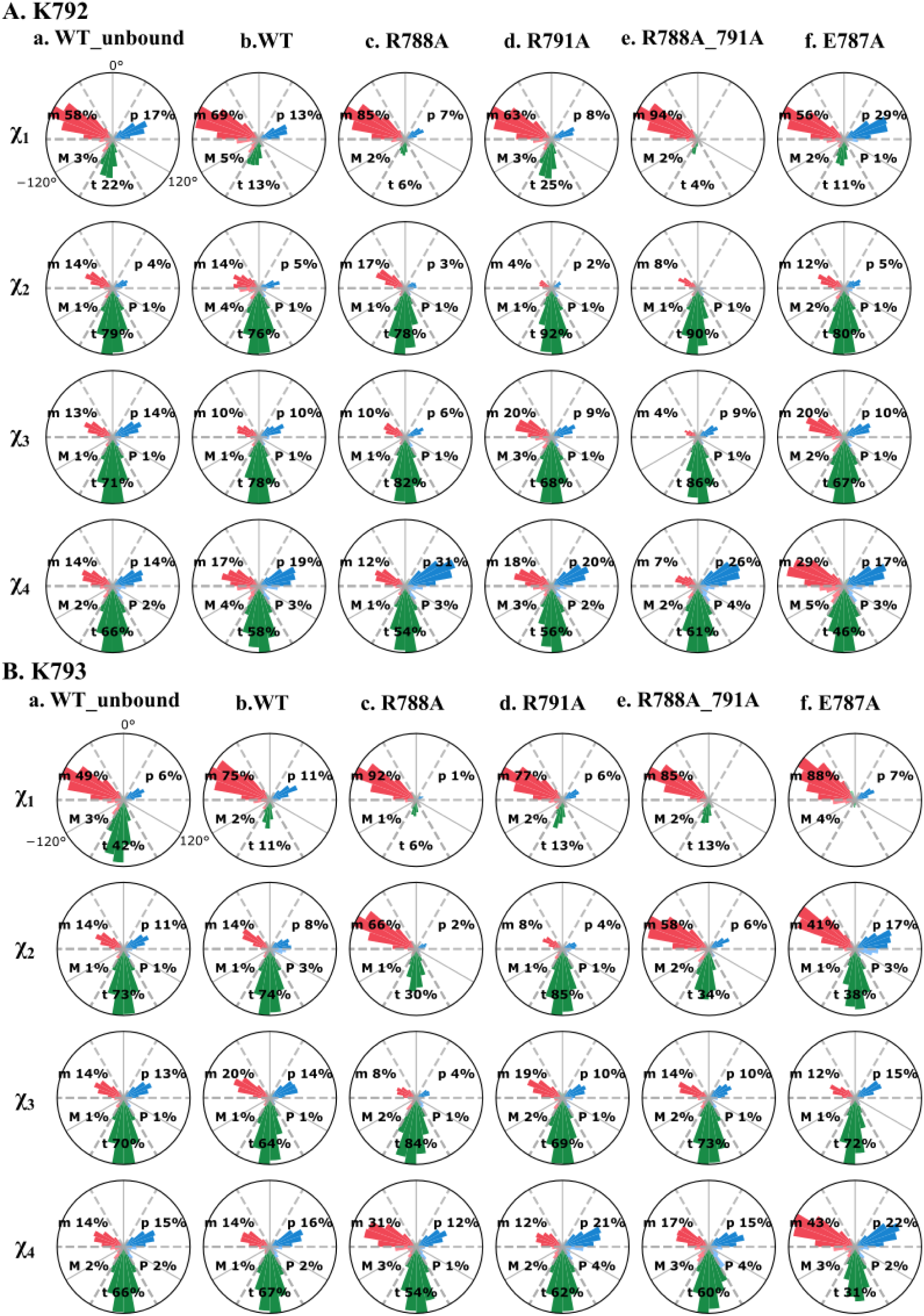
Population distribution of side chain rotamers for Lys792 and Lys793. Polar plots illustrate the distributions of side chain dihedral angles (χ₁-χ₄) for (A) Lys792 and (B) Lys793 across different systems: (a) unbound wild type (WT_unbound), (b) WT, (c) R788A, (d) R791A, (e) double mutant R788A_R791A, and (f) E787A.

**Figure 7.**
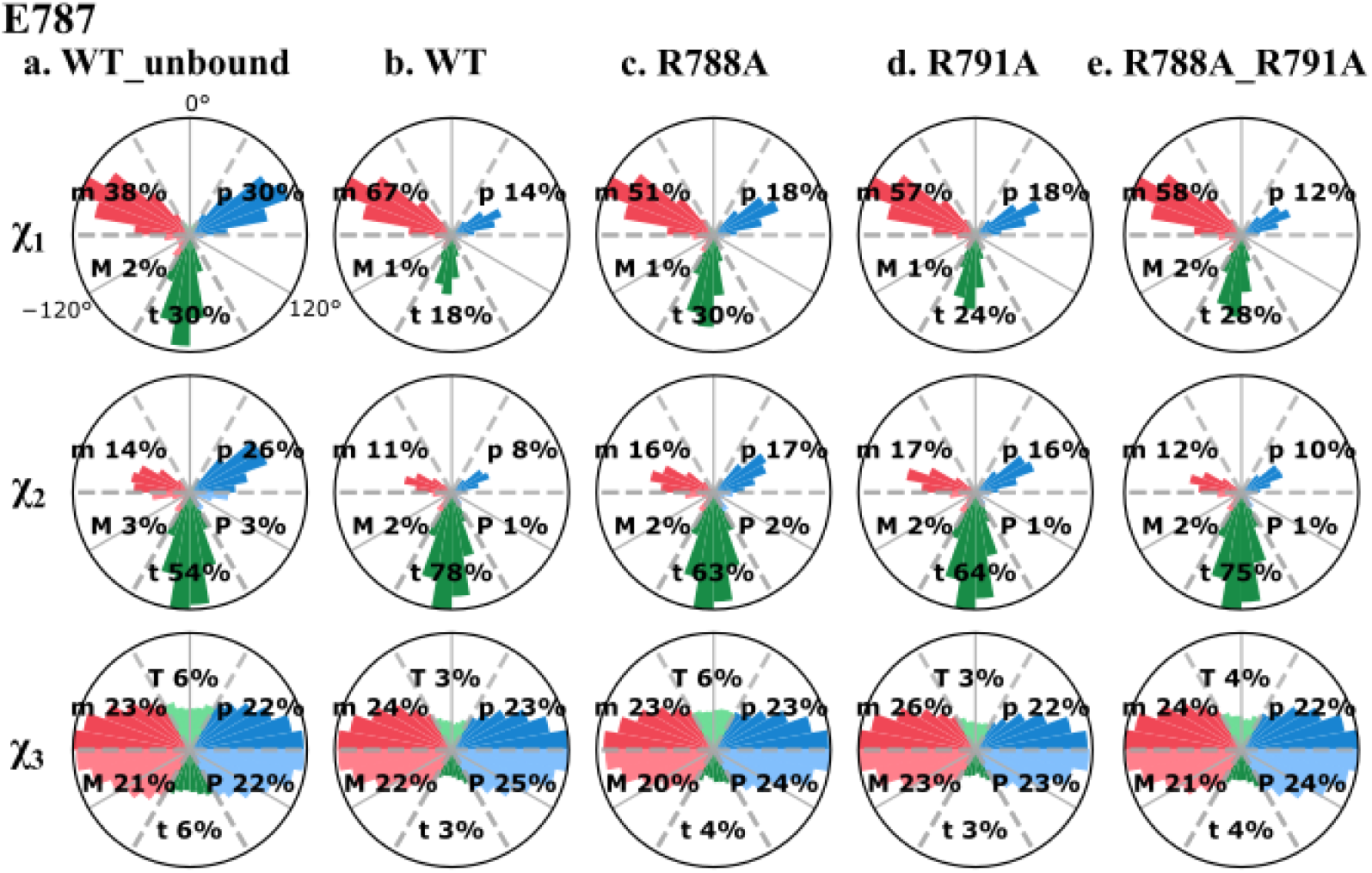
Population distribution of side chain rotamers for Glu787. Polar plots represent the conformational distributions of side chain dihedral angles (χ₁-χ₃) for Glu787 across different systems: (a) unbound wild type (WT_unbound), (b) WT, (c) R788A, (d) R791A, and (e) double mutant R788A_R791A. The distributions are expressed as percentage populations of distinct rotameric states sampled during the simulations. Comparative analysis reveals mutation-dependent shifts in rotamer populations and changes in side chain flexibility of Glu787 upon complex formation.

A comparison of side chain torsions of R788 reveals a complete rearrangement of the rotameric preferences in SF3A1 upon binding SL4 RNA (Figure 5A). χ_1_ mostly occupies *m* state (52%) followed by *t* (23%) and *p* (22%) in WT-unbound. However, 89% of the conformers prefer *p* state in WT. In R791A, *p* and *m* states are equally preferred. On the other hand, *p* state dominates with a population of 77% in E787A. χ_2_ of R788 in WT_unbound shows a strong preference of *t* state (85%) along with a small population distributed in *m*, *p*, *M* and *P* states (Figure 5A). However, the preference of χ_2_ completely shifts to *t* state in WT and two other mutants (R791A and E787A). χ_3_ prefers *t* state (46%) followed by *m* (32%) and *p* (19%) states in WT-unbound. In WT and mutants, the population of *m* state drastically decreases resulting in higher population of *p* and *t* states. In WT and E787A, *p* state population is higher than *t*, whereas p and t states are equally probable in R791A. χ_4_ has an interesting distribution; here off rotameric *M* (18%) and *P* (18%) states are significantly populated along with a higher population of *t* (49%) state. In WT and mutants (R791A and E787A), this distribution is shifted to a combination of on- (*p*) and off- (*P*) rotamers. In WT and E787A, population of *P* state is higher than that of *p,* whereas these two states are equally distributed in R791. The *T* state of χ_5_ in WT_unbound is converted to *t* in WT and mutants.

The side chain torsions of R791 in WT_unbound follows a similar distribution like R788 with minute differences (Figure 5B). However, the rotameric distributions in WT and mutants (R788A and E787A) are quite different from those of R788. In WT, m state of χ_1_ encompasses 90% of the conformations, whereas in R788A, *p* (39%), *m* (25%) and *t* (36%) states are almost evenly populated. In E787A, *p* (72%) state has the highest population followed by *m* (17%) and *t* (11%) states. χ_2_ prefers mostly *m* (59%) and *t* (35%) states in WT. Upon mutation, there is a major population shift from m to *t* state (90% *t* in R788A and 86% *t* in E787A). Conformations of χ_3_ are distributed among different rotameric states in WT. Both the g^+^ state (*p* and *P*) contains 30% population followed by *t* (25%) and *m* (15%) states. In R788A, *P* state is absent while *t* (39%) and *p* (36%) states are almost equally distributed with a less preferred *m* (23%) state.

In E787A, 80% of the conformations are in *p* state. χ_4_ has interesting rotameric preferences and is quite different from R788. In WT, *M* (52%) is he most preferred followed by *t* (23%), P (13%) and *p* (11%). In R788A, the majority of χ_4_ prefers *M* (70%) state along with small preferences of *t* (20%) and *m* (10%) states. Interestingly, 80% of the conformations are in *t* state in E787A. Similar to WT_unbound, χ_5_ remains in *T* state in WT and mutants. In summary, the two Arg residues (R788 and R791), part of the important RGGR motif, sample a wide range of rotameric states. This helps in adjustment of side chain-mediated polar interactions to stabilize the disordered C-terminal in contact with the major groove of the SL4 RNA, while the important interactions are compromised upon mutation.

Unlike Arg, two Lys residues (K792 and K793) present at the tip of the C-terminal have similar rotameric distributions (Figure 6). For example, χ_1_ of K792 highly prefers *m* state, whereas χ_2_, χ_3_ and χ_4_ prefer *t* state (Figure 6A and Table S2). In K793, the rotameric distribution follows a similar trend with a few exceptions. In R788A, R788A_R791A and E787A, χ_2_ of K792 prefers *m* state over *t*, whereas χ_2_ of K793 shows preference for *t* state over *m* (Figure 6B).

The rotamer distribution of E787 in unbound and SL4-RNA-bound states shows some differences (Figure 7). Here, χ1 in WT_unbound almost equally prefers *p* (30%), *m* (38%) and *t* (30%) states. However, in WT and mutants, χ1 highly prefers *m* state (∼50%) followed by *t* (∼25%) and *p* (∼15%) states. On the other hand, χ_2_ prefers *t* state over other rotamers both in unbound and bound states including mutants. Being sp2 hybridized, the side chain of carboxyl carbon of glutamate prefers a planar arrangement, allowing χ_3_ to attain equally probable g+ and g− rotamers. This leads to an almost even population of on- (*p* and *m*) and off- (*P* and *M*) rotamers.

### Conformational analysis of snRNA bound to SF3A1

To examine conformational preference of SL4-RNA in binding SF3A1 and to evaluate the impact of mutations in ULD on overall protein-RNA recognition, we analyzed the RNA conformers in the final 100 ns of all the trajectories. The six torsional angles (α, β, γ, δ, ε, ζ) are projected on pseudo-rotation angles η and θ (Figure 8, Table S3). This pseudo torsion plot provides a reduced representation of the conformational space of a stretch of three nucleotides. Compared to other nucleotides, U12 of UUCG tetraloop shows a significantly wider distribution with multiple clusters. In WT, two clusters are observed: one dense cluster with η: 60° to 120° and θ: 0° to 60°, another sparse cluster with η: 30° to 180° and θ: 180° to 300°. In mutants, the clusters are shifted towards η (ranging from 300° to 360°). The dual mutant, R788A_R791A shows broad distribution with a major cluster shifted towards higher θ. U11, part of the tetraloop, also exhibits significant change in pseudo torsion distribution upon mutation. In WT, U11 sampled a wider range of η (180° to 300°) and θ (180° to 240°). However, this torsional space is restricted with a narrower η (140° to 200°) in E787A. G14 shows unique position of the clusters with θ ranging from 0° to 120°. The terminal nucleotides (G2 and C23) exhibit higher flexibility and the distribution shifts along η upon mutation. C23 is clustered near η: 180° to 240° and θ:180° to 240° in WT and mutants except in R791A. In R791A, the major cluster is shifted towards η:250° to 360° with θ remaining in the same range.

**Figure 8.**
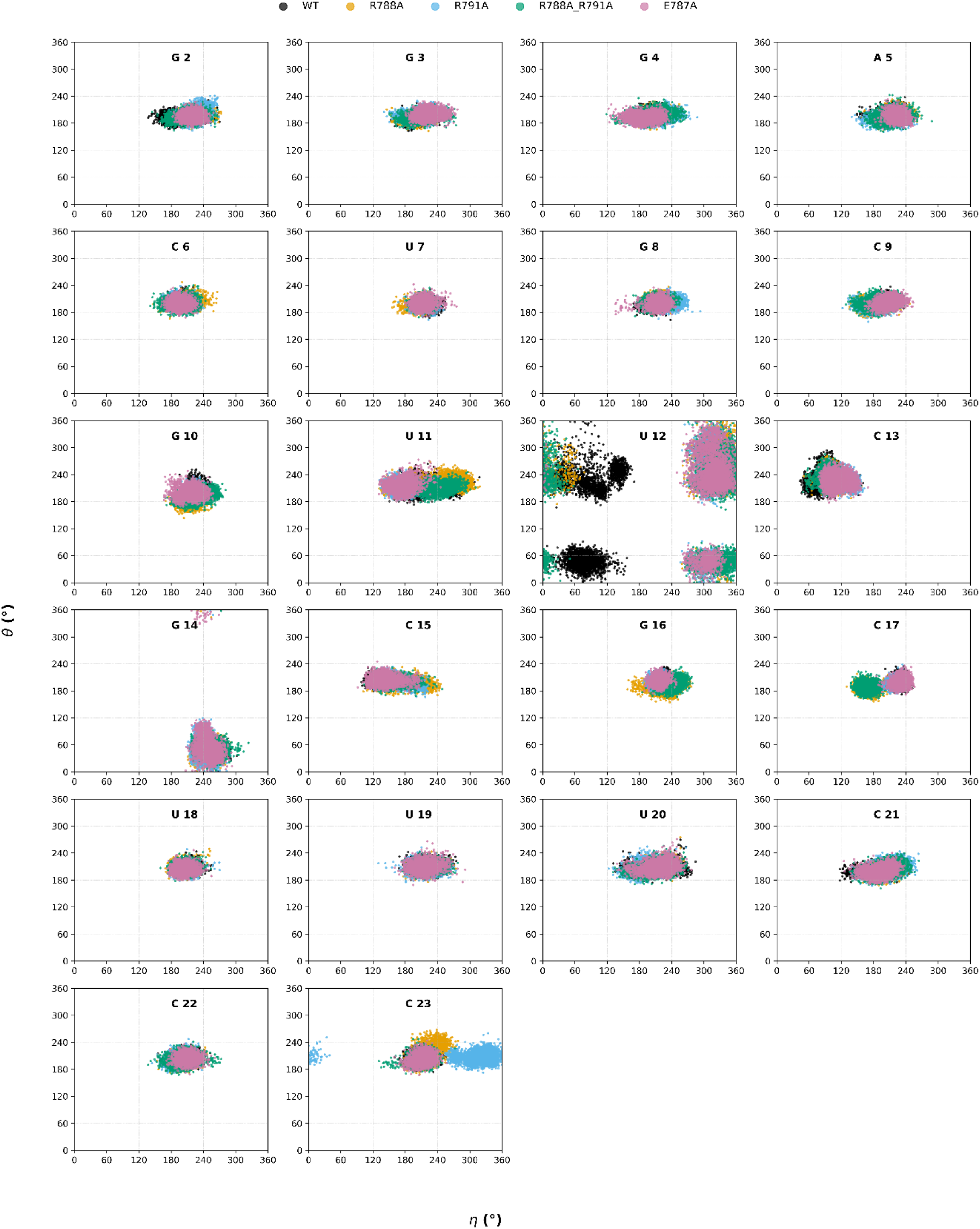
η/θ pseudo-rotation plots for different nucleotides of SL4-snRNA.

We have analyzed the occupancy of backbone and base torsions to get further insights into their dynamics. Occupancy frequencies of backbone (α, β, γ, δ, ε, and ζ) and base (χ) torsion angles for important nucleotides are shown in Figures S3 and S4, with their changes highlighted in black outline. The four nucleotides G10, U11, C13 and C15 contribute significantly to the binding energy across all systems (Refer to Fig. 2A). In contrast, the contributions of U7 and G8 are markedly affected upon mutation, along with C6 and C9. Due to the different impacts of mutation on these two sets of nucleotides, we examine their sugar-phosphate backbone and base torsional angles.

The base torsion distributions differ distinctly between the two groups. For nucleotides C6 to C9, the torsion angles exhibit a narrow distribution, characterized by predominantly low or high occupancy states (Figure S3). There are very few bins with moderate occupancy. In contrast, the second set of nucleotides (G10, U11, C13 and C15) displays broader distributions with moderate occupancy values (Figure S4). Furthermore, the torsional distributions of nucleotides C6 to C9 remain largely unaffected by mutations with a few notable exceptions. In R788A, all four nucleotides exhibit deviations from WT. Specifically, alterations are observed in ε and ζ of C6; β, ε and ζ of U7 and α, β, γ and χ of C9 (Figure S3). In comparison, the second nucleotide set shows more pronounced alterations in their torsional distributions upon mutation. For instance, the χ angle of U11 is evenly distributed between 120° and 240° in WT, whereas in mutants it is predominantly cantered around ∼150°. Other torsion angles of U11 also significantly redistributed upon mutation. Major rearrangements are also observed for ε of G10, and ε and χ of C13 in mutants.

## Discussion

RNA-binding proteins often undergo conformational rearrangement upon binding with partner RNA. They frequently exhibit high sequence specificity and affinity.^37^ However, the non-canonical RNA binding domains often interact with RNA with low affinity through structural motifs rather than sequence-specific manner.^38^ Mutations in these proteins frequently modify their conformational stability and dynamics, resulting in functional changes often associated with pathological conditions.^39^ ULD of SF3A1 typically induces significant structural alterations in both protein and RNA conformations, stabilizing their association. This study investigates the effect of mutually induced conformational changes in the binding of SF3A1 with SL4-snRNA. In addition, we have also explored the effect of mutations in ULD of SF3A1 on RNA binding.

### Non-canonical RNA binding domain and binding affinity

Wild type ULD of SF3A1 binds SL4-RNA with a K_d_ of 160 nM, corresponding to a binding affinity of −9.27 kcal/mol^22^. Mutation of R788 to alanine (R788A) drastically weakens the interaction, increasing the K_d_ and reducing the binding energy to −4.82 kcal/mol. Similarly, R791A mutation results in a moderate loss of affinity, leading to binding energy of −7.10 kcal/mol. To understand the effect of conformational changes in protein and RNA caused by these mutations, we calculate the binding affinity and quantify the contribution of binding pocket residues towards the binding affinity (Figure 3).

Although the absolute values of binding energy are different from that reported in literature, the order of binding affinity follows the experimental trend. Residue-wise decomposition of binding free energy reveals critical energetic contributions from specific interface residues. For example, K786 and E787, located at the C-terminal region of the ULD, exhibit unfavourable energy contributions across all the systems (Figure 3Ai). Whereas, interface Lys (K754, K765, K788, R791, K792 and K793) contribute significantly to the binding affinity. A cooperative behaviour is observed among the residues in R^788^G^789^G^790^R^791^ motif in their contribution to binding energy. Mutation of any of this arginine reduces the contribution of the other one along with the reduction in contribution from the two intermediate glycine residues. This highlights the importance of both the arginine in RGGR motif and emphasizes its importance in ULD-SnRNA recognition.

Interestingly, mutation of E787 to alanine (E787A) shows a minor effect on binding affinity (Kd ∼150 nM, ΔG = −9.30 kcal/mol)^22^. MMGBSA binding energy calculation also results in similar observations where the difference in binding energy between the WT (−112.29 kcal/mol) and E787A (−111.13 kcal/mol) is negligible. This can be explained by persistent backbone mediated H-bonded interactions of E787 with C15 in all the complexes (Figure 3B). Mutations at R788, R791 and K792 clearly demonstrate the crucial role of these basic residues in mediating strong, stabilizing interactions at the interface. Their substitution with alanine eliminated favourable electrostatic and H-bond interactions, resulting in markedly reduced binding affinities and loss of conformational stability. These residues act in a coordinated manner to position and anchor the RNA correctly within the ULD interface. Furthermore, analysis of F763A mutant reveals a significantly reduced binding affinity (K_d_ ∼870 nM, ΔG = −8.26 kcal/mol), which is caused by the disruption of π-π stacking interactions between the aromatic ring of Phe763 and C13 base in UUCG tetraloop^9,22^. This substitution likely alters the kinetics of dissociation by weakening base-stacking interactions that contribute to complex stability. Notably, Phe763 maintains a significant contribution to RNA binding, even in all three mutants, reinforcing its role in stabilizing RNA through aromatic stacking interactions, particularly with C13.

### Side chain rotamer dynamics in protein-RNA recognition

A comparison between side chain torsion of SF3A1 in unbound and bound states reveals that some specific rotameric states are preferred upon complex formation. The difference between rotameric preferences of same amino acids in different systems and different amino acids in the same system can be explained by their surroundings in the complex and the type of interactions they are involved in. R788 In different bound systems (WT and mutants), R788 samples similar rotameric states of the torsions χ_1_ to χ_5_ whereas for R791, preferred rotameric states are changed upon mutation. This can be attributed to the position of the two Arginine residues along the C-terminal axis. Being closely packed into the major groove of snRNA, R788 predominantly interacts with G8 that remain preserved even upon mutations in other neighbouring residues (Figure 3B). This leads to specific rotameric preferences, which are highly preserved across the system (Figure 5A). On the other hand, the side chain of R791 is oriented outwards of the end of the duplex surface. This enables higher flexibility of the R791side chain, resulting in multiple H-bonded interactions with the nucleotides of both the strands. At the 5′ end, R791 interacts with G2, G3 and G4 and at the 3′ end it is involved in interaction with C17 (Figure 3B). These interactions are preserved in WT and mutants with minor variations in their contributions. To interact simultaneously with the nucleotides form both the strands, R791 utilizes multiple side chain rotameric states to preserve structural integrity compensating mutational effects. This has been reflected in broader rotameric distribution across the systems (Figure 5B). Restricted rotameric preferences have been observed for both the lysine residues (K792 and K793) across the systems. This can be attributed to their preference of side chain mediated H-bonds with the nucleotides of the duplex region. In all the systems, both the Lys form a single H-bond in major parts of the simulation time (Figure 4B), leading to similar rotameric states (t state) for χ_2_, χ_3_ and χ_4_ (Figure 6). E787 forms only one backbone mediated H-bond with C15 in 80 to 90% of the simulation time in all systems (Figure 4A). The side chain is not involved in any polar interaction as it is oriented outwards from the major groove of the snRNA. This leads to a similar rotameric preference in all systems (Figure 7). These findings are consistent with the previous studies showing that side chain rotamer stabilization is important for the formation and stability of protein complexes, and mutations at or near interfaces can lead to increased conformational entropy and reduced binding.^40^ Overall, the findings highlight the critical role of rotameric preferences in mediating the structural and functional integrity of the SF3A1-SL4-snRNA complex.

### Conformational dynamics of SL4 RNA in binding SF3A1

The nucleotides of SL4-SnRNA interact with both the globular core of ULD and the unstructured C-terminal of SF3A1. The stem-loop region (G10 to C15) interacts with the globular part, while the duplex region interacts with the C-terminal. The stem-loop region interacts through the UUCG structural motif, whereas the interaction of the duplex region is modulated by the RGGR sequence motif. Mutations impact the interactions of these two regions differently as evident from binding energy decomposition. (Figure 3A(ii)). Two sets of nucleotides from these two regions are chosen for further conformational analyses in terms of their backbone and base torsions. The distribution pattern of torsions is different in the two different sets (Figure S3 and S4). The differences between the two nucleotide sets can be attributed to their distinct structure and interaction. The nucleotides in the duplex region (C6 to C9) form strong base-pair interactions with nucleotides of the opposite strand (G16 to U19), while their backbones interact with amino acid residues. This structural constraint results in narrow torsional distributions and limited sensitivity to mutation. In contrast, the nucleotides G10, U11, C13 and C15, located within the stem loop region, exhibits weaker base-pairing but stronger protein-RNA interactions, particularly with residues preceding the RGGR motif (G10-K786, U11-K765, C13-K756 and C15-E787). These interactions remain largely preserved upon mutation, as supported by lifetime analysis of interface H-bonds (Figure 3B). It allows these nucleotides to sample a wider range of torsional conformations, enabling them to maintain strong interactions with interface residues even when mutations perturb other regions of the complex.

In each mutant, the weakening or loss of favourable nucleotide interactions coincides with the corresponding disruption of the adjacent protein-RNA contacts. These findings collectively underscore the cooperative relationship between protein interface residues and specific nucleotides, highlighting the mutations at critical protein sites propagate through the interface to diminish the energetic contributions of nearby nucleotides, thereby weakening overall binding affinity.

## Conclusions

Dysregulation of splicing machinery, often stemming from impaired molecular interactions, is implicated in numerous human diseases. This study elucidates the structural role of non-canonical binding of ULD domain of SF3A1 with U1 snRNA to mediate early spliceosomal assembly. The conformational dynamics of interface residues and nucleotides reveal a mutual recognition mechanism driven by the RGGR motif of SF3A1 and the UUCG tetraloop of SL4-snRNA. The C-terminal RGGR motif recognizes the duplex region of SL4-snRNA. On the other hand, nucleotides in the tetraloop region of SL4-snRNA are involved in interactions with the globular region of the ULD domain. Mutational studies highlight the effect of mutation on these recognition processes. Upon mutation of residues (R788 and R791) of RGGR motif, a substantial loss in protein-RNA interaction is observed as reflected in binding energy values. E787 shows a persistent backbone-mediated interaction with nucleotides and its mutation does not affect the overall binding energy. Due to stronger base pairing, nucleotides of the stem region show less conformational variability upon mutation. On the other hand, nucleotides of the UUCG tetraloop significantly adopt their conformations to maintain stable interaction with the globular part of the protein to maintain overall structural integrity.

## Acknowledgement

SK, MB and SM acknowledge Indian Institute of Technology Kharagpur for their fellowship and infrastructure support. AM and RPB acknowledge DB, Govt. of India for computational facility funded by the BIC programme of DBT, Govt. of India (Gran no. BT/PR40175/BTIS/137/41/2022). The authors acknowledge the Param Shakti Supercomputing facility at Indian Institute of Technology Kharagpur.

## Notes

### Competing Interest Statement

The authors have declared no competing interest.

## References

(1) Jurica, M. S.; Moore, M. J. Pre-mRNA Splicing: Awash in a Sea of Proteins. Molecular Cell 2003, 12 (1), 5–14. 10.1016/S1097-2765(03)00270-3.

(2) Kastner, B.; Will, C. L.; Stark, H.; Lührmann, R. Structural Insights into Nuclear Pre-mRNA Splicing in Higher Eukaryotes. Cold Spring Harb Perspect Biol 2019, 11 (11), a032417. 10.1101/cshperspect.a032417.

(3) Rogalska, M. E.; Vivori, C.; Valcárcel, J. Regulation of Pre-mRNA Splicing: Roles in Physiology and Disease, and Therapeutic Prospects. Nature Reviews Genetics 2023, 24 (4), 251–269. 10.1038/s41576-022-00556-8.

(4) Fica, S. M.; Oubridge, C.; Galej, W. P.; Wilkinson, M. E.; Bai, X. C.; Newman, A. J.; Nagai, K. Structure of a Spliceosome Remodelled for Exon Ligation. Nature 2017, 542 (7641), 377–380. 10.1038/NATURE21078.

(5) Sperling, J.; Azubel, M.; Sperling, R. Structure and Function of the Pre-mRNA Splicing Machine. Structure 2008, 16 (11), 1605–1615. 10.1016/J.STR.2008.08.011.

(6) Zhan, X.; Lu, Y.; Shi, Y. Molecular Basis for the Activation of Human Spliceosome. Nat Commun 2024, 15 (1), 6348. 10.1038/s41467-024-50785-0.

(7) Martelly, W.; Fellows, B.; Kang, P.; Vashisht, A.; Wohlschlegel, J. A.; Sharma, S. Synergistic Roles for Human U1 snRNA Stem-Loops in Pre-mRNA Splicing. RNA Biology 2021, 18 (12), 2576–2593. 10.1080/15476286.2021.1932360.

(8) Martelly, W.; Fellows, B.; Senior, K.; Marlowe, T.; Sharma, S. Identification of a Noncanonical RNA Binding Domain in the U2 snRNP Protein SF3A1. RNA 2019, 25 (11), 1509–1521. 10.1261/rna.072256.119.

(9) De Vries, T.; Martelly, W.; Campagne, S.; Sabath, K.; Sarnowski, C. P.; Wong, J.; Leitner, A.; Jonas, S.; Sharma, S.; Allain, F. H. T. Sequence-Specific RNA Recognition by an RGG Motif Connects U1 and U2 snRNP for Spliceosome Assembly. Proceedings of the National Academy of Sciences of the United States of America 2022, 119 (6). 10.1073/pnas.2114092119.

(10) Kuwasako, K.; He, F.; Inoue, M.; Tanaka, A.; Sugano, S.; Güntert, P.; Muto, Y.; Yokoyama, S. Solution Structures of the SURP Domains and the Subunit-Assembly Mechanism within the Splicing Factor SF3a Complex in 17S U2 snRNP. Structure 2006, 14 (11), 1677–1689. 10.1016/J.STR.2006.09.009.

(11) Zhang; Yan, C.; Zhan, X.; Li, L.; Lei, J.; Shi, Y. Structure of the Human Activated Spliceosome in Three Conformational States. Cell Research 2018, 28 (3), 307–322. 10.1038/CR.2018.14.

(12) Krämer, A.; Mulhauser, F.; Wersig, C.; Gröning, K.; Bilbe, G. Mammalian Splicing Factor SF3a120 Represents a New Member of the SURP Family of Proteins and Is Homologous to the Essential Splicing Factor PRP21p of Saccharomyces Cerevisiae. RNA 1995, 1 (3), 260–272.

(13) Hochstrasser, M. Origin and Function of Ubiquitin-like Proteins. Nature 2009, 458 (7237), 422–429. 10.1038/NATURE07958.

(14) Jentsch, S.; Pyrowolakis, G. Ubiquitin and Its Kin: How Close Are the Family Ties? Trends in Cell Biology 2000, 10 (8), 335–342. 10.1016/S0962-8924(00)01785-2.

(15) Liu, Z.; Luyten, I.; Bottomley, M. J.; Messias, A. C.; Houngninou-Molango, S.; Sprangers, R.; Zanier, K.; Krämer, A.; Sattler, M. Structural Basis for Recognition of the Intron Branch Site RNA by Splicing Factor 1. Science 2001, 294 (5544), 1098–1102. 10.1126/SCIENCE.1064719.

(16) Winget, J. M.; Mayor, T. The Diversity of Ubiquitin Recognition: Hot Spots and Varied Specificity. Molecular Cell 2010, 38 (5), 627–635. 10.1016/J.MOLCEL.2010.05.003.

(17) Martínez-Lumbreras, S.; Morguet, C.; Sattler, M. Dynamic Interactions Drive Early Spliceosome Assembly. Current Opinion in Structural Biology 2024, 88, 102907. 10.1016/j.sbi.2024.102907.

(18) Wahl, M. C.; Will, C. L.; Lührmann, R. The Spliceosome: Design Principles of a Dynamic RNP Machine. Cell 2009, 136 (4), 701–718. 10.1016/J.CELL.2009.02.009.

(19) Cieśla, M.; Ngoc, P. C. T.; Cordero, E.; Martinez, Á. S.; Morsing, M.; Muthukumar, S.; Beneventi, G.; Madej, M.; Munita, R.; Jönsson, T.; Lövgren, K.; Ebbesson, A.; Nodin, B.; Hedenfalk, I.; Jirström, K.; Vallon-Christersson, J.; Honeth, G.; Staaf, J.; Incarnato, D.; Pietras, K.; Bosch, A.; Bellodi, C. Oncogenic Translation Directs Spliceosome Dynamics Revealing an Integral Role for SF3A3 in Breast Cancer. Molecular Cell 2021, 81 (7), 1453–1468.e12. 10.1016/j.molcel.2021.01.034.

(20) Visconte, V.; Makishima, H.; Maciejewski, J. P.; Tiu, R. V. Emerging Roles of the Spliceosomal Machinery in Myelodysplastic Syndromes and Other Hematological Disorders. Leukemia 2012, 26 (12), 2447–2454. 10.1038/leu.2012.130.

(21) Scotti, M. M.; Swanson, M. S. RNA Mis-Splicing in Disease. Nat Rev Genet 2016, 17 (1), 19–32. 10.1038/nrg.2015.3.

(22) Nameki, N.; Terawaki, S. I.; Takizawa, M.; Kitamura, M.; Muto, Y.; Kuwasako, K. Structural Insights into Recognition of SL4, the UUCG Stem-Loop, of Human U1 snRNA by the Ubiquitin-like Domain, Including the C-Terminal Tail in the SF3A1 Subunit of U2 snRNP. The Journal of Biochemistry 2023, 174 (2), 203–216. 10.1093/JB/MVAD033.

(23) Molinaro, M.; Tinoco, I. Use of Ultra Stable UNCG Tetraloop Hairpins to Fold RNA Structures: Thermodynamic and Spectroscopic Applications. Nucleic Acids Research 1995, 23 (15), 3056–3063. 10.1093/NAR/23.15.3056.

(24) Berman, H. M.; Westbrook, J.; Feng, Z.; Gilliland, G.; Bhat, T. N.; Weissig, H.; Shindyalov, I. N.; Bourne, P. E. The Protein Data Bank. Nucleic acids research 2000, 28 (1), 235–242. 10.1093/NAR/28.1.235.

(25) Pettersen, E. F.; Goddard, T. D.; Huang, C. C.; Couch, G. S.; Greenblatt, D. M.; Meng, E. C.; Ferrin, T. E. UCSF Chimera-A Visualization System for Exploratory Research and Analysis. J Comput Chem 2004, 25 (13), 1605–1612. 10.1002/jcc.20084.

(26) Mark, P.; Nilsson, L. Structure and Dynamics of the TIP3P, SPC, and SPC/E Water Models at 298 K. Journal of Physical Chemistry A 2001, 105 (43), 9954–9960. 10.1021/jp003020w.

(27) Götz, A. W.; Williamson, M. J.; Xu, D.; Poole, D.; Le Grand, S.; Walker, R. C. Routine Microsecond Molecular Dynamics Simulations with AMBER on GPUs. 1. Generalized Born. Journal of Chemical Theory and Computation 2012, 8 (5), 1542–1555. 10.1021/CT200909J.

(28) Bauer, P.; Hess, B.; Lindahl, E. GROMACS 2022 Source Code, 2022. 10.5281/ZENODO.6103835.

(29) Bussi, G.; Donadio, D.; Parrinello, M. Canonical Sampling through Velocity Rescaling. The Journal of Chemical Physics 2007, 126 (1), 014101. 10.1063/1.2408420.

(30) Parrinello, M.; Rahman, A. Polymorphic Transitions in Single Crystals: A New Molecular Dynamics Method. Journal of Applied Physics 1981, 52 (12), 7182–7190. 10.1063/1.328693.

(31) Hess, B. P-LINCS: A Parallel Linear Constraint Solver for Molecular Simulation. Journal of Chemical Theory and Computation 2008, 4 (1), 116–122. 10.1021/ct700200b.

(32) Darden, T.; York, D.; Pedersen, L. Particle Mesh Ewald: An N⋅log(N) Method for Ewald Sums in Large Systems. The Journal of Chemical Physics 1993, 98 (12), 10089–10092. 10.1063/1.464397.

(33) Roe, D. R.; Cheatham, T. E. PTRAJ and CPPTRAJ: Software for Processing and Analysis of Molecular Dynamics Trajectory Data. Journal of Chemical Theory and Computation 2013, 9 (7), 3084–3095. 10.1021/ct400341p.

(34) Valdés-Tresanco, M. S.; Valdés-Tresanco, M. E.; Valiente, P. A.; Moreno, E. gmx_MMPBSA: A New Tool to Perform End-State Free Energy Calculations with GROMACS. J. Chem. Theory Comput. 2021, 17 (10), 6281–6291. 10.1021/acs.jctc.1c00645.

(35) McGibbon, R. T.; Beauchamp, K. A.; Harrigan, M. P.; Klein, C.; Swails, J. M.; Hernández, C. X.; Schwantes, C. R.; Wang, L.-P.; Lane, T. J.; Pande, V. S. MDTraj: A Modern Open Library for the Analysis of Molecular Dynamics Trajectories. Biophysical Journal 2015, 109 (8), 1528–1532. 10.1016/j.bpj.2015.08.015.

(36) Bottaro, S.; Bussi, G.; Pinamonti, G.; Reißer, S.; Boomsma, W.; Lindorff-Larsen, K. Barnaba: Software for Analysis of Nucleic Acid Structures and Trajectories. RNA 2019, 25 (2), 219–231. 10.1261/rna.067678.118.

(37) Glisovic, T.; Bachorik, J. L.; Yong, J.; Dreyfuss, G. RNA-Binding Proteins and Post-Transcriptional Gene Regulation. FEBS Letters 2008, 582 (14), 1977–1986. 10.1016/J.FEBSLET.2008.03.004.

(38) Ray, D.; Laverty, K. U.; Jolma, A.; Nie, K.; Samson, R.; Pour, S. E.; Tam, C. L.; Von Krosigk, N.; Nabeel-Shah, S.; Albu, M.; Zheng, H.; Perron, G.; Lee, H.; Najafabadi, H.; Blencowe, B.; Greenblatt, J.; Morris, Q.; Hughes, T. R. RNA-Binding Proteins That Lack Canonical RNA-Binding Domains Are Rarely Sequence-Specific. Sci Rep 2023, 13 (1). 10.1038/s41598-023-32245-9.

(39) Stefl, S.; Nishi, H.; Petukh, M.; Panchenko, A. R.; Alexov, E. Molecular Mechanisms of Disease-Causing Missense Mutations. Journal of Molecular Biology 2013, 425 (21), 3919–3936. 10.1016/j.jmb.2013.07.014.

(40) Kortemme, T.; Baker, D. A Simple Physical Model for Binding Energy Hot Spots in Protein–Protein Complexes. Proc. Natl. Acad. Sci. U.S.A. 2002, 99 (22), 14116–14121. 10.1073/pnas.202485799.

