## Supporting Information for "Decoding Mutually Induced Conformational Changes in Non-Canonical Recognition of U1 SL4 snRNA by ULD of SF3A1 during Early Spliceosome Assembly"

**for**

Ph: +91-3222-283790

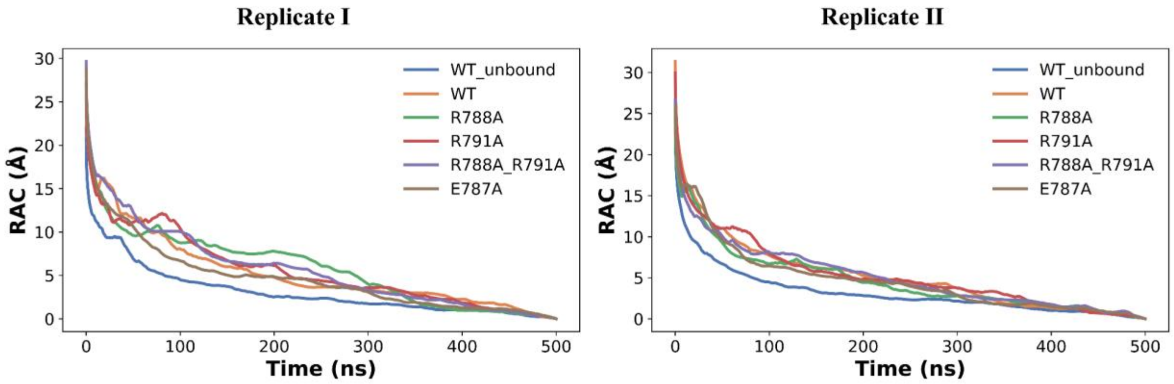


Figure S1. Root-mean-square (RMS) average correlation (RAC) curves for all the simulated systems in both the independent replicate simulations.

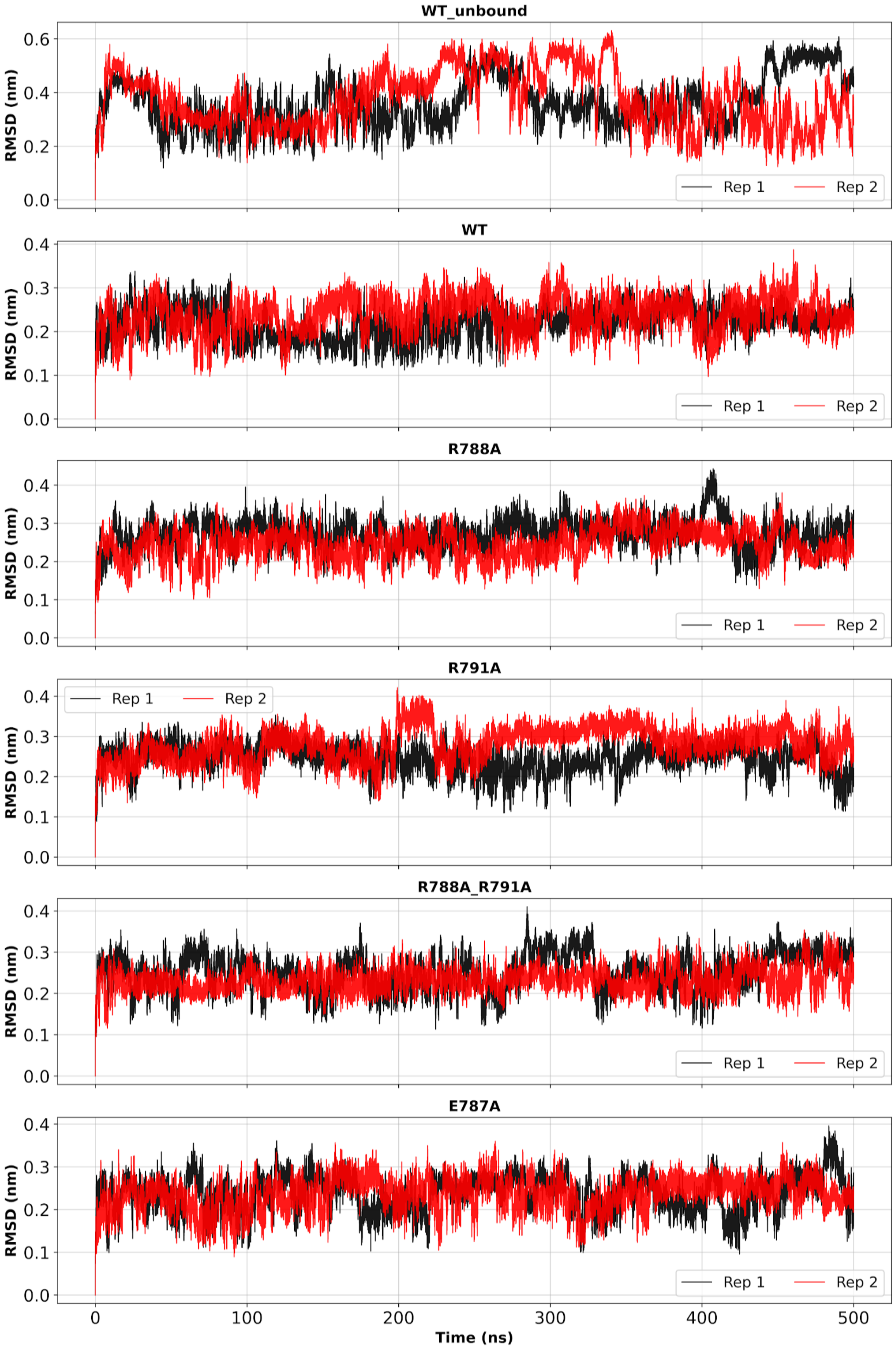


Figure S2A. RMSD profile for all the simulated systems for both the independent replicates.

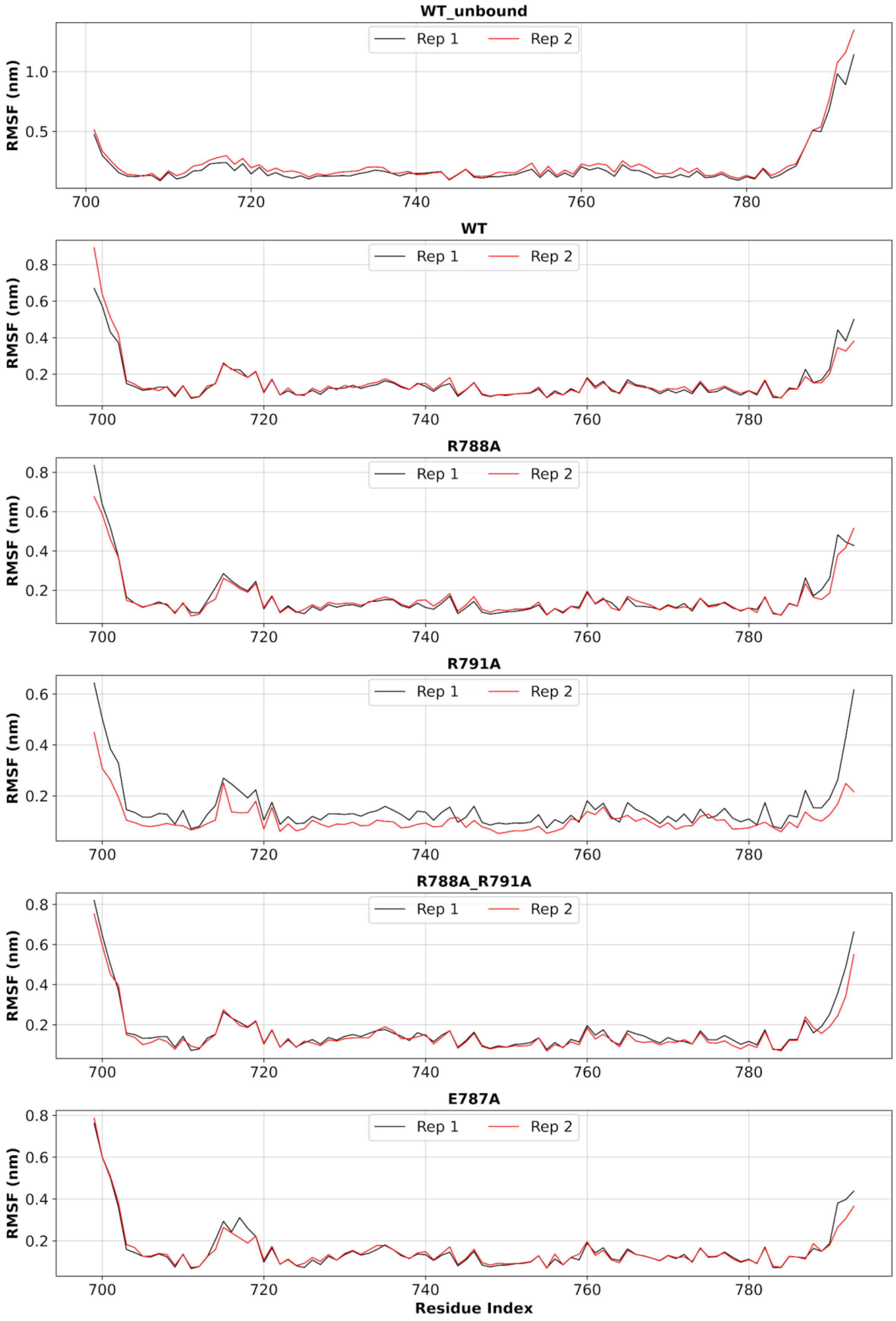


Figure S2B. RMSF profile for all simulated systems for both the independent replicate.

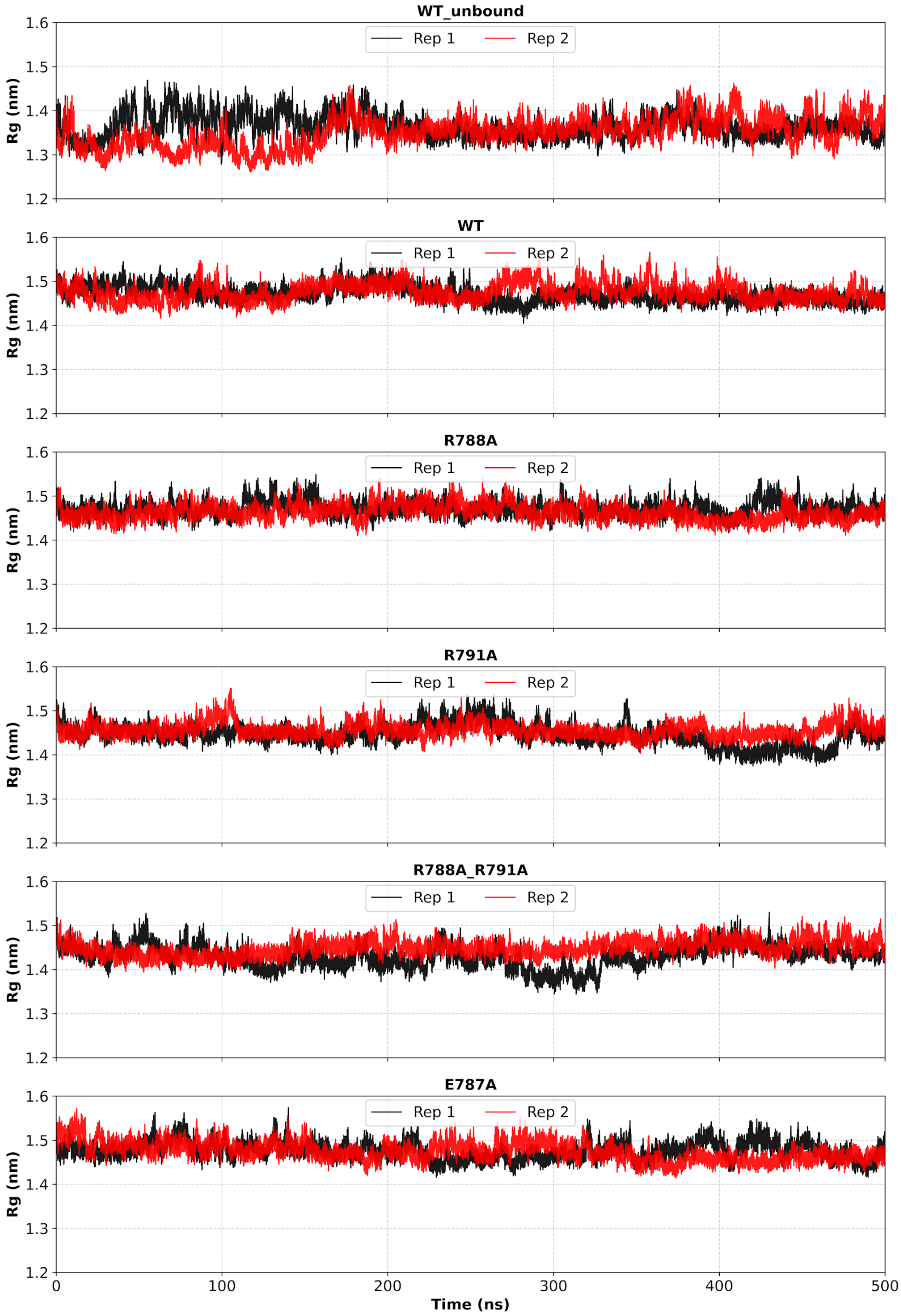


Figure S2C. Rg profile for all simulated systems for both the independent replicates.

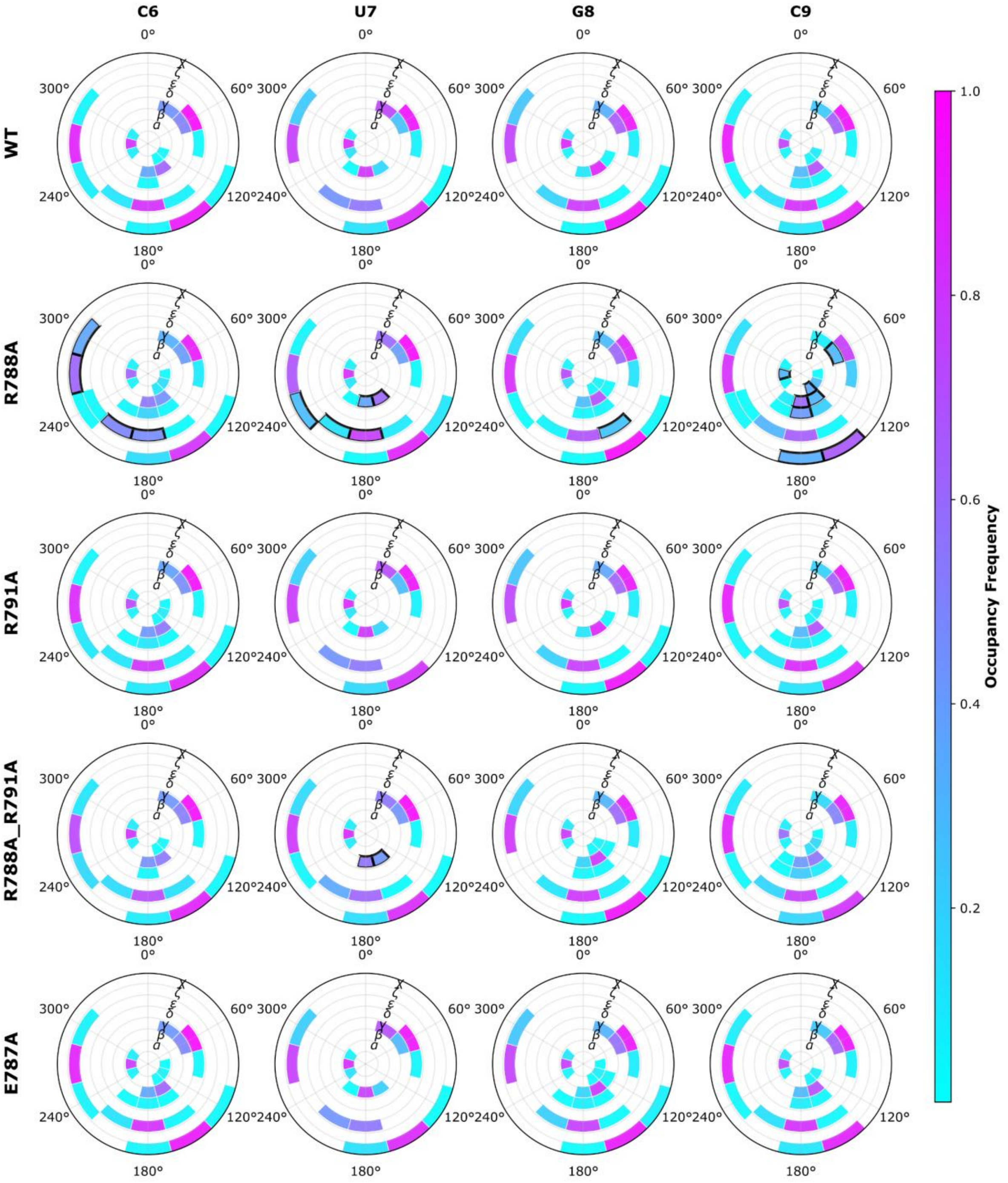


Figure S3. Wheel plots showing the occupancy frequencies of RNA backbone torsion angles (α, β, γ, δ, ε, ζ) and glycosidic χ angle for C6, U7, G8, and C9 nucleotides in the SL4-snRNA during last 100 ns of simulation. Each wheel represents one nucleotide, with coloured sectors indicating conformational populations across the full 0° to 360° range. Intensity of colour represents the occupancy of rotamers.

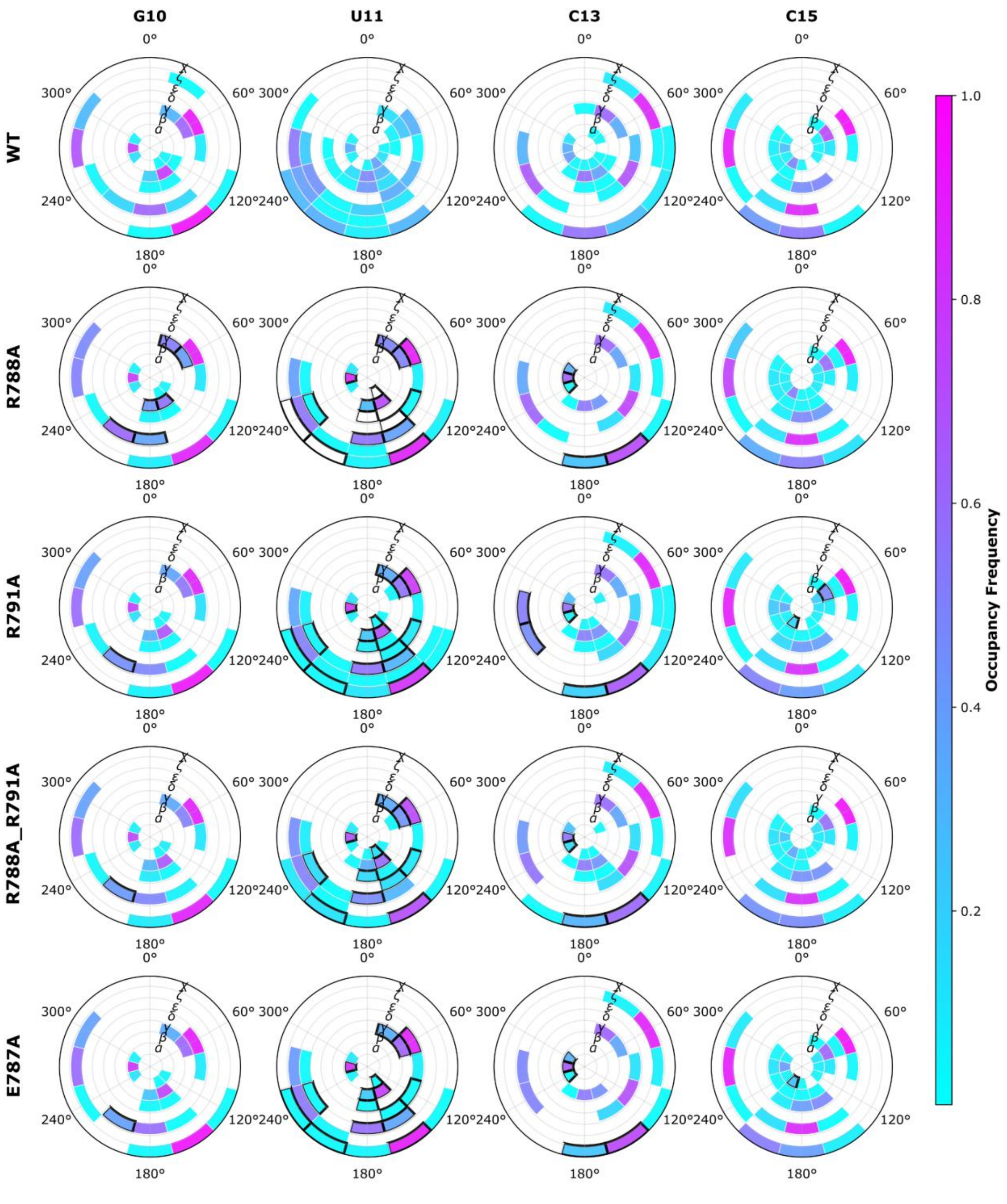


Figure S4. Wheel plots showing the occupancy frequencies of RNA backbone torsion angles (α, β, γ, δ, ε, ζ) and glycosidic χ angle for the G10, U11, C13 and C15 tetraloop nucleotides of SL4-snRNA during the last 100 ns of simulations. Each wheel represents one nucleotide, with coloured sectors indicating conformational populations across the full 0°-360° range.

Table S1. Rotamer distribution of Glu 787, Arg 788 and Arg 791 in the ULD of SF3A1.

| systems | Side chain torsion angle | On-rotamer | | | Off-rotamer | | |
|---|---|---|---|---|---|---|---|
| | | %*p* | %*t* | %*m* | %*P* | %*T* | %*M* |
| | | | ***Glu 787*** | | | | |
| WT_unbound | χ1 | 30.3 | 33.4 | 34.2 | 0.2 | 0.3 | 1.6 |
| WT_unbound | χ2 | 26.1 | 56.6 | 12.9 | 2 | 0 | 2.3 |
| WT_unbound | χ3 | 21.6 | 5.7 | 24.2 | 22.3 | 5.4 | 20.8 |
| WT | χ1 | 16.3 | 17.1 | 65.3 | 0.2 | 0.1 | 1.1 |
| WT | χ2 | 6.5 | 76.6 | 14.1 | 0.7 | 0 | 2.1 |
| WT | χ3 | 22.8 | 3.4 | 24.3 | 25.3 | 3.1 | 21.1 |
| R788A | χ1 | 26.8 | 22.8 | 48.8 | 0.2 | 0.1 | 1.3 |
| R788A | χ2 | 11.6 | 60 | 24.7 | 1.3 | 0 | 2.4 |
| R788A | χ3 | 22.9 | 3.8 | 23.1 | 24.6 | 5.7 | 20 |
| R791A | χ1 | 16.6 | 19.5 | 62.3 | 0.2 | 0.1 | 1.3 |
| R791A | χ2 | 13.7 | 66.5 | 17 | 0.7 | 0 | 2.2 |
| R791A | χ3 | 21.7 | 3 | 26 | 23 | 3.1 | 23.2 |
| R788A_R791A | χ1 | 9.9 | 25.1 | 63.1 | 0.1 | 0.1 | 1.6 |
| R788A_R791A | χ2 | 7.1 | 79.2 | 10.7 | 0.8 | 0 | 2.1 |
| R788A_R791A | χ3 | 23.2 | 3.6 | 23.3 | 25.2 | 3.5 | 21.3 |
| | | | ***Arg 788*** | | | | |
| WT_unbound | χ1 | 20.7 | 24.1 | 51.8 | 0.3 | 0.1 | 3 |
| WT_unbound | χ2 | 6.3 | 85.9 | 4.9 | 1.6 | 0 | 1.4 |
| WT_unbound | χ3 | 19.1 | 42.9 | 35 | 1.3 | 0.1 | 1.6 |
| WT_unbound | χ4 | 6.6 | 49.8 | 7.1 | 18.8 | 0 | 17.8 |
| WT_unbound | χ5 | 0.4 | 0 | 0.3 | 0 | 99.2 | 0 |
| WT | χ1 | 74.8 | 7.5 | 17.2 | 0.3 | 0 | 0.2 |
| WT | χ2 | 0 | 99.9 | 0 | 0.1 | 0 | 0 |
| WT | χ3 | 74.9 | 17.3 | 7.3 | 0.1 | 0 | 0.2 |
| WT | χ4 | 55.2 | 0 | 0 | 44.7 | 0 | 0 |
| WT | χ5 | 0 | 99.8 | 0 | 0.2 | 0 | 0 |
| R791A | χ1 | 64.8 | 1 | 33.9 | 0.2 | 0 | 0.1 |
| R791A | χ2 | 0 | 99.6 | 0 | 0.2 | 0 | 0.1 |
| R791A | χ3 | 64.8 | 33.9 | 1 | 0.2 | 0 | 0.1 |
| R791A | χ4 | 52 | 0.1 | 0 | 47.8 | 0 | 0 |
| R791A | χ5 | 0 | 99.8 | 0 | 0.2 | 0 | 0 |
| E787A | χ1 | 67.3 | 7.7 | 24.6 | 0.2 | 0 | 0.2 |
| E787A | χ2 | 0.1 | 99.2 | 0.1 | 0.3 | 0 | 0.4 |
| E787A | χ3 | 67.2 | 24.6 | 7.6 | 0.2 | 0 | 0.4 |
| E787A | χ4 | 49 | 3.1 | 0 | 47.8 | 0 | 0 |
| E787A | χ5 | 0 | 99.8 | 0 | 0.2 | 0 | 0.1 |
| | | | ***Arg 791*** | | | | |

| systems | Side chain torsion angle | On-rotamer | | | Off-rotamer | | |
|---|---|---|---|---|---|---|---|
| | | %*p* | %*t* | %*m* | %*P* | %*T* | %*M* |
| WT_unbound | χ1 | 13.1 | 26.5 | 55.7 | 0.2 | 0.3 | 4.3 |
| WT_unbound | χ2 | 7.8 | 81.5 | 8.1 | 1.3 | 0 | 1.3 |
| WT_unbound | χ3 | 23.8 | 52.4 | 20.1 | 1.6 | 0.1 | 1.9 |
| WT_unbound | χ4 | 8.7 | 45.9 | 7.8 | 19.4 | 0 | 18.2 |
| WT_unbound | χ5 | 0.4 | 0 | 0.4 | 0 | 99.2 | 0 |
| WT | χ1 | 8.6 | 3.1 | 87.2 | 0.1 | 0.1 | 0.9 |
| WT | χ2 | 2.9 | 36.5 | 59.5 | 0.6 | 0 | 0.5 |
| WT | χ3 | 31.8 | 21.1 | 10.3 | 35.6 | 0 | 1.1 |
| WT | χ4 | 5.5 | 30.4 | 1.7 | 10.2 | 0 | 52.3 |
| WT | χ5 | 0.2 | 0 | 0.4 | 0 | 99.4 | 0 |
| R788A | χ1 | 18.1 | 20.9 | 58.3 | 0.3 | 0.1 | 2.3 |
| R788A | χ2 | 2.5 | 69.1 | 26.1 | 1.1 | 0 | 1.2 |
| R788A | χ3 | 28.6 | 51.2 | 16.3 | 2.2 | 0.1 | 1.6 |
| R788A | χ4 | 6.2 | 51 | 8.5 | 12.1 | 0 | 22.2 |
| R788A | χ5 | 0.3 | 0 | 0.5 | 0 | 99.1 | 0 |
| E787A | χ1 | 16.1 | 31 | 48.5 | 0.1 | 0.1 | 4.1 |
| E787A | χ2 | 22.4 | 48.9 | 26.1 | 1.2 | 0.2 | 1.2 |
| E787A | χ3 | 42.5 | 23 | 26.6 | 6.8 | 0.2 | 0.9 |
| E787A | χ4 | 6.5 | 50.6 | 2.7 | 16 | 0 | 24.2 |
| E787A | χ5 | 0.4 | 0 | 0.5 | 0 | 99.2 | 0 |

Table S2. Rotamer distribution of C-terminal Lys residues in ULD of SF3A1.

| systems | Side-chain torsion angle | On-rotamer | | | Off-rotamer | | |
|---|---|---|---|---|---|---|---|
| | | %*p* | %*t* | %*m* | %*P* | %*T* | %*M* |
| | | Lys 786 | | | | | |
| WT_unbound | χ1 | 18.1 | 54.7 | 20.7 | 0.3 | 0 | 6.2 |
| WT_unbound | χ2 | 3.1 | 92.3 | 1.9 | 1.2 | 0 | 1.4 |
| WT_unbound | χ3 | 16.8 | 70.9 | 9.7 | 1.6 | 0.1 | 1 |
| WT_unbound | χ4 | 10.8 | 75.7 | 9 | 2.3 | 0 | 2.2 |
| WT | χ1 | 0.4 | 91.8 | 6 | 0.1 | 0 | 1.7 |
| WT | χ2 | 9.3 | 85.8 | 1.6 | 1.5 | 0 | 1.7 |
| WT | χ3 | 4.9 | 79.9 | 14 | 0.6 | 0 | 0.5 |
| WT | χ4 | 22.4 | 66.9 | 5.1 | 4 | 0 | 1.5 |
| R788A | χ1 | 4 | 85.8 | 5.4 | 0.1 | 0 | 4.7 |
| R788A | χ2 | 4.4 | 90 | 0.9 | 3.5 | 0 | 1.3 |
| R788A | χ3 | 7.2 | 87.3 | 4 | 1 | 0 | 0.4 |
| R788A | χ4 | 23.9 | 63.1 | 6.2 | 5.6 | 0.1 | 1.1 |
| R791A | χ1 | 1 | 94.4 | 2.3 | 0.1 | 0 | 2.2 |
| R791A | χ2 | 15.6 | 74.8 | 5 | 1.7 | 0.1 | 2.8 |
| R791A | χ3 | 0.8 | 96.1 | 1.8 | 0.7 | 0 | 0.6 |
| R791A | χ4 | 24.2 | 58.7 | 10.9 | 3.9 | 0.1 | 2.1 |
| R788A_R791A | χ1 | 0.2 | 93.8 | 3.5 | 0.1 | 0 | 2.3 |
| R788A_R791A | χ2 | 12.4 | 81.3 | 2.7 | 1.7 | 0 | 1.8 |
| R788A_R791A | χ3 | 2.8 | 94.3 | 1.7 | 0.8 | 0 | 0.4 |
| R788A_R791A | χ4 | 20.5 | 61.8 | 11.5 | 3.6 | 0 | 2.5 |
| E787A | χ1 | 0.2 | 91 | 6.3 | 0 | 0 | 2.3 |
| E787A | χ2 | 10.8 | 83 | 2.8 | 1.8 | 0 | 1.6 |
| E787A | χ3 | 3.4 | 92.6 | 2.7 | 0.8 | 0 | 0.6 |
| E787A | χ4 | 20.5 | 64.6 | 8.4 | 4.6 | 0 | 1.9 |
| | | Lys 792 | | | | | |
| WT_unbound | χ1 | 17.4 | 17 | 62.6 | 0.4 | 0.1 | 2.5 |
| WT_unbound | χ2 | 3.8 | 78.1 | 15.5 | 1.1 | 0 | 1.4 |
| WT_unbound | χ3 | 14 | 70.2 | 13.6 | 1 | 0.1 | 1.1 |
| WT_unbound | χ4 | 14.5 | 66.3 | 14.3 | 2.6 | 0 | 2.1 |
| WT | χ1 | 10.8 | 18.9 | 66.3 | 0.2 | 0 | 3.8 |
| WT | χ2 | 7.7 | 72.5 | 15.4 | 1.3 | 0.1 | 3 |
| WT | χ3 | 11.1 | 75.6 | 11.2 | 1.1 | 0.1 | 1 |
| WT | χ4 | 17.2 | 57.6 | 18.5 | 2.9 | 0 | 3.8 |
| R788A | χ1 | 19.7 | 23.3 | 54.1 | 0.3 | 0.1 | 2.5 |
| R788A | χ2 | 4 | 78.8 | 14.7 | 1.3 | 0 | 1.2 |
| R788A | χ3 | 10.9 | 74.3 | 12.7 | 1 | 0.1 | 1 |
| R788A | χ4 | 15 | 65.3 | 15.1 | 2.1 | 0.1 | 2.4 |
| R791A | χ1 | 13.3 | 26.7 | 56.3 | 0.3 | 0.1 | 3.3 |
| R791A | χ2 | 7.5 | 76.7 | 13.2 | 1.1 | 0.1 | 1.5 |

| systems | Side-chain torsion angle | On-rotamer | | | Off-rotamer | | |
|---|---|---|---|---|---|---|---|
| | | %*p* | %*t* | %*m* | %*P* | %*T* | %*M* |
| R791A | χ3 | 11.3 | 72.7 | 13.4 | 1.1 | 0.1 | 1.4 |
| R791A | χ4 | 23.4 | 51.3 | 19.8 | 2.9 | 0.1 | 2.5 |
| R788A_R791A | χ1 | 4.8 | 17 | 74.3 | 0.2 | 0.1 | 3.7 |
| R788A_R791A | χ2 | 6.5 | 78.4 | 12.6 | 1.3 | 0 | 1.2 |
| R788A_R791A | χ3 | 15.4 | 70.8 | 11.4 | 1.1 | 0.1 | 1.2 |
| R788A_R791A | χ4 | 23.2 | 57.8 | 14.2 | 2.7 | 0.1 | 1.9 |
| E787A | χ1 | 18 | 10.8 | 67.3 | 0.2 | 0.1 | 3.7 |
| E787A | χ2 | 6.4 | 81.1 | 9.7 | 1.7 | 0.1 | 1 |
| E787A | χ3 | 8 | 66.4 | 22.5 | 0.7 | 0.1 | 2.3 |
| E787A | χ4 | 19.7 | 46 | 27.8 | 2.8 | 0.1 | 3.8 |
| | | | | Lys 793 | | | |
| WT_unbound | χ1 | 6.2 | 40.9 | 49.7 | 0.1 | 0.1 | 2.9 |
| WT_unbound | χ2 | 11.2 | 72.8 | 13.6 | 1.2 | 0 | 1.2 |
| WT_unbound | χ3 | 13.2 | 70.4 | 13.8 | 1.1 | 0.1 | 1.4 |
| WT_unbound | χ4 | 16.1 | 65.1 | 14.2 | 2.2 | 0 | 2.5 |
| WT | χ1 | 7.7 | 12 | 78.1 | 0.1 | 0.2 | 1.9 |
| WT | χ2 | 8.3 | 69.4 | 17.9 | 3.6 | 0.1 | 0.6 |
| WT | χ3 | 9.6 | 73.7 | 14.7 | 0.7 | 0.1 | 1.2 |
| WT | χ4 | 17.2 | 63.5 | 15.6 | 2 | 0.1 | 1.7 |
| R788A | χ1 | 2.8 | 7 | 87.9 | 0 | 0.1 | 2.2 |
| R788A | χ2 | 3.6 | 54.6 | 39.5 | 1.3 | 0.1 | 0.9 |
| R788A | χ3 | 5.7 | 75.8 | 16.2 | 0.9 | 0.1 | 1.4 |
| R788A | χ4 | 15.4 | 57.3 | 21.6 | 2.9 | 0 | 2.8 |
| R791A | χ1 | 12.8 | 20.4 | 63.9 | 0.1 | 0.2 | 2.6 |
| R791A | χ2 | 10.1 | 72.7 | 12.7 | 3 | 0 | 1.5 |
| R791A | χ3 | 14.9 | 65.7 | 17.2 | 1 | 0.1 | 1.2 |
| R791A | χ4 | 17.2 | 64 | 14.2 | 2.5 | 0 | 2 |
| R788A_R791A | χ1 | 3.1 | 20.4 | 74.3 | 0 | 0.1 | 2.1 |
| R788A_R791A | χ2 | 8.3 | 50 | 39.6 | 0.8 | 0 | 1.4 |
| R788A_R791A | χ3 | 14.1 | 69 | 14.5 | 1.4 | 0.1 | 0.8 |
| R788A_R791A | χ4 | 11 | 57.2 | 25.6 | 2.6 | 0 | 3.6 |
| E787A | χ1 | 12.3 | 11.9 | 73.2 | 0.1 | 0.2 | 2.3 |
| E787A | χ2 | 11.6 | 62.4 | 22.1 | 3 | 0.1 | 0.9 |
| E787A | χ3 | 12.5 | 70.6 | 14.7 | 0.9 | 0.1 | 1.2 |
| E787A | χ4 | 20.5 | 51.7 | 23.4 | 2 | 0.1 | 2.2 |

Table S3. Values of average pseudo torsion angle for the 22 nucleotides in different systems

| nucleotide | η | | θ | |
|---|---|---|---|---|
| | mean | Std. dev | mean | Std. dev |
| WT | | | | |
| G_2 | 201.68 | 24.77 | 195.37 | 9.21 |
| G_3 | 205.88 | 14.76 | 194.09 | 8.12 |
| G_4 | 207.57 | 11.24 | 198.57 | 7.50 |
| A_5 | 219.81 | 11.73 | 200.72 | 8.66 |
| C_6 | 200.56 | 8.51 | 203.80 | 8.19 |
| U_7 | 219.99 | 9.62 | 195.29 | 7.54 |
| G_8 | 216.38 | 9.36 | 197.59 | 7.42 |
| C_9 | 203.20 | 8.38 | 203.20 | 7.43 |
| G_10 | 228.61 | 9.08 | 204.46 | 9.38 |
| U_11 | 216.07 | 29.84 | 210.64 | 11.92 |
| U_12 | 83.92 | 34.28 | 114.69 | 92.06 |
| C_13 | 92.74 | 16.93 | 228.24 | 16.08 |
| G_14 | 254.94 | 9.46 | 46.37 | 10.84 |
| C_15 | 134.18 | 10.24 | 203.60 | 7.42 |
| G_16 | 220.25 | 6.98 | 206.29 | 8.03 |
| C_17 | 227.63 | 7.33 | 199.80 | 8.44 |
| U_18 | 211.22 | 9.69 | 206.41 | 7.74 |
| U_19 | 220.11 | 14.16 | 208.82 | 8.49 |
| U_20 | 217.17 | 23.03 | 206.23 | 8.88 |
| C_21 | 191.97 | 18.73 | 199.23 | 9.44 |

| nucleotide | η | | θ | |
|---|---|---|---|---|
| | mean | Std. dev | mean | Std. dev |
| C_22 | 208.87 | 13.20 | 202.53 | 7.98 |
| C_23 | 211.07 | 11.39 | 205.38 | 8.60 |
| R788A | | | | |
| G_2 | 220.24 | 12.71 | 198.56 | 7.84 |
| G_3 | 204.04 | 15.08 | 194.05 | 8.12 |
| G_4 | 210.53 | 11.97 | 197.64 | 8.15 |
| A_5 | 221.34 | 12.55 | 197.20 | 8.91 |
| C_6 | 207.87 | 13.17 | 201.42 | 8.90 |
| U_7 | 211.97 | 10.18 | 199.21 | 7.73 |
| G_8 | 216.95 | 12.79 | 203.40 | 7.79 |
| C_9 | 191.62 | 17.56 | 198.95 | 8.36 |
| G_10 | 214.03 | 12.42 | 184.44 | 11.72 |
| U_11 | 239.83 | 50.97 | 220.04 | 11.82 |
| U_12 | 306.25 | 82.53 | 239.25 | 52.51 |
| C_13 | 101.29 | 11.95 | 230.73 | 13.95 |
| G_14 | 250.49 | 13.34 | 44.28 | 13.63 |
| C_15 | 164.57 | 27.15 | 199.34 | 8.69 |
| G_16 | 230.16 | 17.47 | 188.90 | 9.60 |
| C_17 | 170.67 | 11.62 | 186.97 | 8.05 |
| U_18 | 199.96 | 10.17 | 206.72 | 7.92 |
| U_19 | 215.97 | 9.85 | 209.30 | 8.09 |

| nucleotide | η | | θ | |
|---|---|---|---|---|
| | mean | Std. dev | mean | Std. dev |
| U_20 | 219.66 | 14.02 | 209.16 | 9.17 |
| C_21 | 200.74 | 17.67 | 198.98 | 9.52 |
| C_22 | 205.76 | 11.91 | 200.50 | 8.72 |
| C_23 | 218.54 | 13.65 | 208.50 | 14.41 |
| R791A | | | | |
| G_2 | 220.46 | 12.76 | 198.48 | 10.51 |
| G_3 | 206.99 | 15.84 | 196.05 | 8.28 |
| G_4 | 217.11 | 15.54 | 196.36 | 8.52 |
| A_5 | 215.46 | 15.89 | 192.30 | 8.36 |
| C_6 | 194.00 | 13.71 | 199.32 | 7.85 |
| U_7 | 209.69 | 8.20 | 197.73 | 7.49 |
| G_8 | 232.88 | 11.63 | 202.50 | 8.26 |
| C_9 | 180.70 | 14.17 | 195.37 | 8.36 |
| G_10 | 225.49 | 8.54 | 200.81 | 8.58 |
| U_11 | 172.86 | 12.59 | 210.37 | 10.15 |
| U_12 | 314.10 | 25.17 | 216.12 | 81.64 |
| C_13 | 119.04 | 16.18 | 226.51 | 11.58 |
| G_14 | 241.33 | 11.45 | 60.10 | 22.75 |
| C_15 | 154.91 | 21.03 | 202.22 | 8.50 |
| G_16 | 211.94 | 7.67 | 200.85 | 7.80 |
| C_17 | 225.28 | 7.97 | 199.31 | 7.79 |

| nucleotide | η | | θ | |
|---|---|---|---|---|
| | mean | Std. dev | mean | Std. dev |
| U_18 | 207.53 | 10.02 | 204.14 | 7.82 |
| U_19 | 214.51 | 14.26 | 204.96 | 8.53 |
| U_20 | 200.84 | 19.40 | 207.26 | 10.43 |
| C_21 | 208.84 | 18.98 | 205.39 | 11.13 |
| C_22 | 201.45 | 13.29 | 202.20 | 10.23 |
| C_23 | 318.60 | 39.26 | 207.34 | 9.64 |
| R788A_R791A | | | | |
| G_2 | 212.47 | 15.18 | 195.11 | 8.18 |
| G_3 | 210.26 | 15.56 | 195.02 | 8.01 |
| G_4 | 211.01 | 16.76 | 198.46 | 8.64 |
| A_5 | 215.54 | 15.92 | 194.97 | 9.53 |
| C_6 | 203.03 | 14.90 | 199.64 | 8.86 |
| U_7 | 214.40 | 8.59 | 199.59 | 7.24 |
| G_8 | 211.12 | 12.95 | 202.20 | 7.89 |
| C_9 | 189.95 | 16.67 | 201.44 | 9.07 |
| G_10 | 229.39 | 16.22 | 195.84 | 10.21 |
| U_11 | 239.44 | 16.97 | 209.64 | 8.80 |
| U_12 | 285.56 | 111.33 | 161.68 | 100.97 |
| C_13 | 95.51 | 10.74 | 231.86 | 14.28 |
| G_14 | 254.89 | 15.50 | 52.67 | 11.01 |
| C_15 | 150.74 | 15.86 | 201.86 | 7.77 |

| nucleotide | η | | θ | |
|---|---|---|---|---|
| | mean | Std. dev | mean | Std. dev |
| G_16 | 245.58 | 9.29 | 195.04 | 8.81 |
| C_17 | 166.96 | 9.37 | 188.10 | 8.36 |
| U_18 | 205.22 | 10.23 | 207.62 | 7.91 |
| U_19 | 218.43 | 13.14 | 209.03 | 8.05 |
| U_20 | 209.99 | 19.19 | 206.20 | 9.50 |
| C_21 | 202.22 | 20.06 | 203.12 | 10.72 |
| C_22 | 206.31 | 14.98 | 201.92 | 9.57 |
| C_23 | 211.86 | 11.72 | 203.34 | 9.03 |
| E787A | | | | |
| G_2 | 219.82 | 10.75 | 196.35 | 7.65 |
| G_3 | 221.70 | 13.79 | 200.28 | 8.16 |
| G_4 | 189.32 | 17.09 | 192.26 | 7.82 |
| A_5 | 226.66 | 9.28 | 195.60 | 8.05 |
| C_6 | 196.85 | 9.96 | 198.88 | 8.20 |
| U_7 | 212.01 | 9.68 | 199.50 | 8.04 |
| G_8 | 215.45 | 10.22 | 199.78 | 8.11 |
| C_9 | 208.13 | 11.22 | 202.14 | 8.10 |
| G_10 | 218.86 | 12.30 | 201.32 | 9.51 |
| U_11 | 176.17 | 12.77 | 213.18 | 11.49 |
| U_12 | 314.37 | 26.82 | 211.44 | 82.85 |
| C_13 | 115.58 | 15.88 | 226.46 | 12.61 |

| nucleotide | η | | θ | |
|---|---|---|---|---|
| | mean | Std. dev | mean | Std. dev |
| G_14 | 248.49 | 11.82 | 58.68 | 31.46 |
| C_15 | 140.35 | 15.22 | 205.17 | 8.40 |
| G_16 | 215.84 | 6.47 | 203.61 | 8.13 |
| C_17 | 231.26 | 7.71 | 201.23 | 8.20 |
| U_18 | 202.48 | 10.41 | 204.29 | 7.89 |
| U_19 | 215.75 | 12.81 | 208.34 | 9.39 |
| U_20 | 218.06 | 17.39 | 208.51 | 9.84 |
| C_21 | 190.07 | 16.30 | 198.30 | 9.62 |
| C_22 | 210.10 | 10.31 | 204.14 | 8.00 |
| C_23 | 214.59 | 10.94 | 203.86 | 9.31 |
